# DDB2 localizes to the nucleolus to mediate oxaliplatin resistance in colorectal cancer

**DOI:** 10.64898/2026.11.23.768665

**Authors:** Vinod Kumar, Ashna Nagpal, Asema Maratova, Vera Roginskaya, Katherine E. Helfrich, Callen T. Wallace, Michael J. Calderon, Simon C. Watkins, Dennis Hsu, Orlando D. Schärer, Bennett Van Houten

**Affiliations:** UPMC Hillman Cancer Center, University of Pittsburgh, Pittsburgh, PA 15213, USA; Department of Pharmacology and Chemical Biology, School of Medicine, University of Pittsburgh, Pittsburgh, PA 15213, USA; Center for Genomic Integrity, Institute for Basic Science, Ulsan, Republic of Korea; Department of Biological Sciences, College of Information-Bio Convergence Engineering, Ulsan National Institute of Science and Technology, Ulsan, Republic of Korea; Center for Biologic Imaging, University of Pittsburgh, PA, USA; Department of Medicine, University of Pittsburgh School of Medicine, Pittsburgh, PA, USA; The Ohio State University Comprehensive Cancer Center – Arthur G. James Cancer Hospital and Richard J. Solove Research Institute (OSUCCC – James), Columbus, OH 43210, USA; Department of Internal Medicine, Division of Medical Oncology, The Ohio State University, Columbus, OH 43210, USA

## Abstract

Despite the widespread use of platinum drugs in cancer therapy, defining their mechanisms of action and predicting clinical outcomes remain challenging. Unlike cisplatin, which kills cells mainly through the DNA damage response, oxaliplatin toxicity is driven largely by nucleolar stress, yet how cells cope with this stress remains poorly understood. A nucleotide excision repair protein, damage-specific DNA binding protein 2 (DDB2), protects cells from oxaliplatin but not cisplatin, suggesting that it may preferentially engage oxaliplatin adducts. Using single-molecule analysis of DNA-binding proteins from nuclear extracts, we show that DDB2 binds both oxaliplatin and cisplatin intrastrand adducts. Oxaliplatin, but not cisplatin, drives progressive accumulation of DDB2 in nucleoli, pointing to a role at sites of oxaliplatin-induced nucleolar stress. The repair-deficient xeroderma pigmentosum group E K244E variant localizes to nucleoli but fails to rescue oxaliplatin sensitivity of DDB2-knockout cells, indicating that repair activity, not nucleolar localization alone, is required for protection. In colorectal cancer (CRC), elevated DDB2 levels correlate with higher oxaliplatin IC_50_, are higher in tumors than adjacent normal tissue by immunohistochemistry, and DDB2 transcript associates with non-response and shorter progression-free survival in FOLFOX-treated patients. Pharmacological inactivation or degradation of DDB2 synergizes with oxaliplatin, offering a biomarker-driven strategy to overcome chemoresistance.

## Introduction

Oxaliplatin administered in combination regimens is a standard-of-care adjuvant therapy for curable colorectal cancer (CRC) and is a mainstay in managing metastatic CRC ^1^. Unfortunately, a substantial proportion of patients receiving adjuvant FOLFOX (5-fluorouracil/leucovorin/oxaliplatin) inevitably experience disease recurrence. Furthermore, oxaliplatin treatment inflicts significant toxicities, including dose-limiting peripheral sensory neuropathy, which can become permanent ^2^. Currently, there are no universally established, or FDA-approved predictive biomarkers for routine clinical use to predict an individual’s response to oxaliplatin. Thus, it would be helpful to determine which patients would benefit most from this drug, develop reliable biomarkers for drug response, and identify new drug targets.

Damage-specific DNA binding protein 2 (DDB2) is the first responder for damage recognition within chromatin during global genome nucleotide excision repair (GG-NER) of cyclobutane pyrimidine dimers (CPDs) and 6-4 photoproducts (6-4PPs) ^3–6^. DDB2, together with DDB1, forms a heterodimeric complex that associates with CUL4A-ROC1, creating an E3 ubiquitin ligase. This repair complex works to detect DNA photolesions and remodel chromatin to allow subsequent repair ^7–11^. Alterations in the DDB2 protein, such as K244E, cause a skin cancer-prone syndrome xeroderma pigmentosum complementation group E (XP-E) ^12^. UV-DDB has a broad damage specificity beyond UV-induced photoproducts, including cisplatin adducts and abasic sites ^13–15^. Our laboratory has shown that in addition to UV damage, UV-DDB recognizes base damage including: 8-oxoG, hypoxanthine, and alkylation damage ^3, 16–19^. Consequently, DDB2 is now recognized as a multifaceted regulator of chromatin remodeling after DNA damage ^20–22^. A recent study ^23^ using DDB2 knockout (KO) cell lines indicated that these cells were sensitive to oxaliplatin, but not cisplatin, and the authors concluded that DDB2 does not efficiently recognize cisplatin adducts in cells. However, this study did not directly quantify the binding affinities of DDB2 to oxaliplatin and cisplatin adducts using biochemical experiments. Earlier reports by Payne et al. ^13, 14^ demonstrated that UV-DDB has specific binding affinity for oligonucleotides containing cisplatin adducts. These discrepancies underscore the importance of reconciling cellular and biochemical studies of nucleotide excision repair in processing platinum-DNA adducts.

The cytotoxicity of platinum drugs is generally thought to arise from Pt-DNA adducts. Previous studies show that oxaliplatin generates less DNA damage than cisplatin but is comparably cytotoxic ^24^, suggesting differences in their mechanisms of toxicity. A study has shown that while initial lesion frequency correlates with cisplatin, carboplatin, and oxaliplatin concentrations, resistant cell lines often exhibit significantly lower initial lesion frequencies ^25^. Other recent work has suggested that cisplatin kills cells by DNA damage response (DDR) and oxaliplatin by inducing stress to the nucleolus ^26, 27^.

Here we investigated the apparent discrepancy that DDB2 KO cells are sensitive to oxaliplatin but not cisplatin. To further understand and clarify this differential response, we examined DDB2 binding to site-specific oxaliplatin and cisplatin DNA adducts. We show that DDB2 recognizes both oxaliplatin and cisplatin adducts but displays higher affinity for oxaliplatin adducts. Furthermore, we discover that treatment with oxaliplatin, but not cisplatin, causes DDB2 accumulation in the nucleolus with mutual colocalization of NPM1. Interestingly, we found that the NER-deficient DDB2 K244E variant that enters the nucleolus fails to protect cells from oxaliplatin. We further demonstrate that elevated DDB2 mRNA levels are significantly associated with non-responder status compared to responder status in CRC patients treated with oxaliplatin/FOLFOX. Since mRNA and protein levels do not always directly correlate, we developed and validated an immunohistochemistry (IHC) assay using DDB2-specific antibodies to verify that DDB2 protein levels are differentially expressed in colon cancer patient tissues. We found that high DDB2-expressing CRC cell lines have higher IC_50_ for oxaliplatin. Finally, we investigated the combination of oxaliplatin with two small-molecule inactivators of DDB2: CSN5i-3 (DDB2 degrader) and NAEi/MLN4924/Pevonedistat (DDB2 inhibitor). We report that the combination of DDB2 depletion/inactivation and oxaliplatin yields synergistic effects. These findings provide a framework for predicting therapeutic efficacy (i.e., low tumor DDB2 levels indicate a likely response) and provide a proof of principle pharmacological approach to treat tumors expressing high levels of DDB2 that would otherwise be resistant to oxaliplatin treatment.

## Results

### DDB2 protein levels vary across colorectal cancer cell lines and correlate with oxaliplatin, but not cisplatin resistance

A recent study ^23^ reports that oxaliplatin resistance in tumor cells depends on DDB2 expression levels, and that NER damage sensor XPC is not recruited to oxaliplatin DNA adducts in cells in the absence of DDB2. To investigate the relationship between DDB2 expression and platinum-drug resistance in colorectal cancer, we studied a panel of CRC cell lines (HCT116, SW480, HT29 and RKO). We first measured basal DDB2 protein levels in these CRC lines alongside U2OS cells and RPE cells (a non-cancerous retinal pigment epithelial line) (Fig. 1a). Western blot analysis revealed substantial variability in DDB2 protein levels across these lines, with HCT116 and SW480 expressing the highest levels and RPE the lowest (Fig. 1b, c). We also examined XPC and p21, two proteins functionally linked to DDB2 and nucleotide excision repair (NER), and found that their expression did not track with DDB2 levels across the panel (Fig. 1d-f), suggesting that DDB2 abundance is not simply a proxy for general NER pathway activity in these cells.

**Fig. 1.**
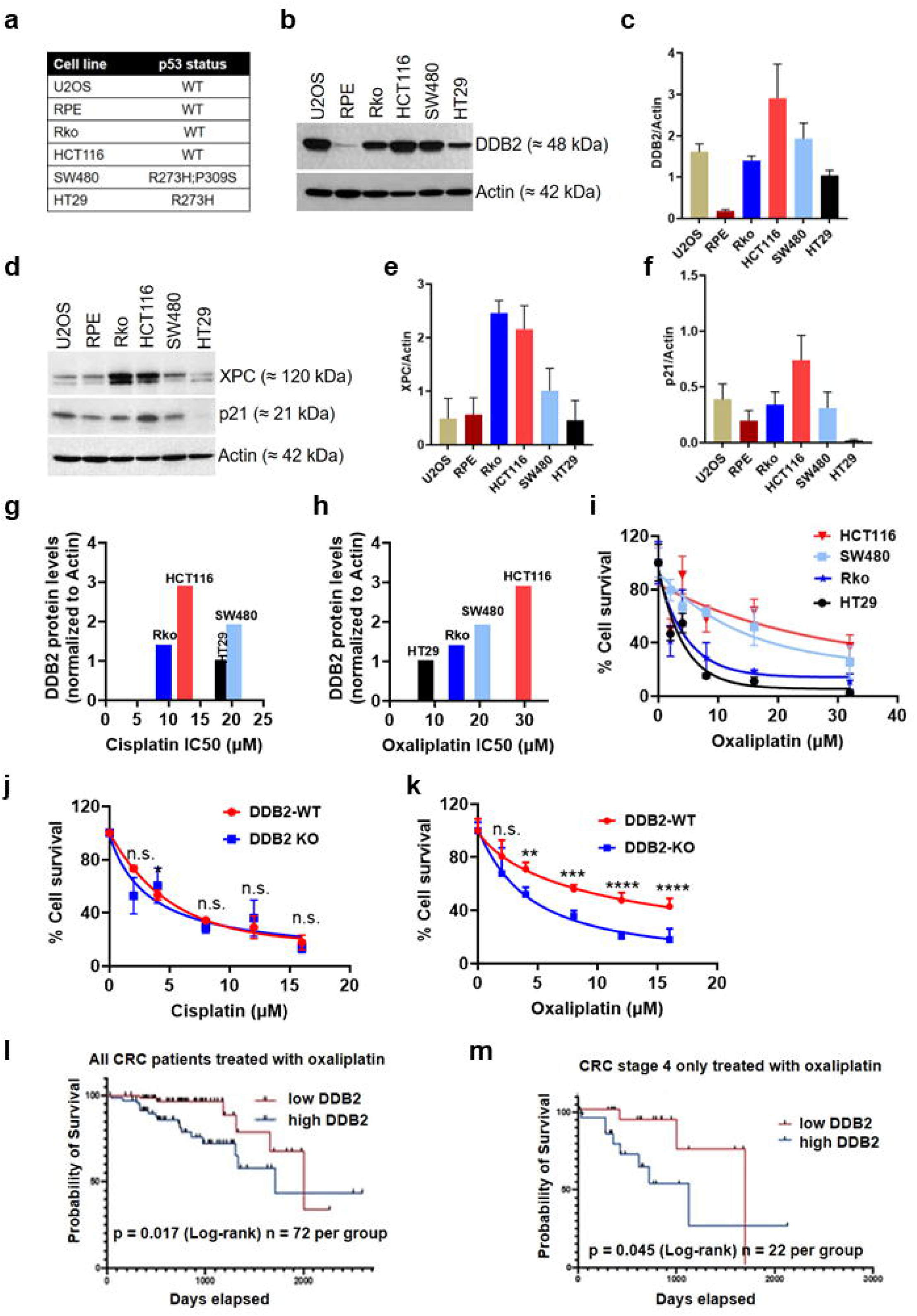
DDB2 protein expression correlates with oxaliplatin, but not cisplatin, resistance in colorectal cancer cell lines and predicts survival in oxaliplatin-treated patients. **a** TP53 mutational status of cell lines used (U2OS, RPE, Rko, HCT116, SW480, HT29). b Representative western blot of DDB2 and actin (loading control) in indicated cell lines. c Quantification of DDB2 levels normalized to actin from b. Mean ± s.e.m.; n = 3 independent exposures. d Representative western blot of XPC and p21, with actin as loading control. Quantification of e XPC and f p21 levels normalized to actin from d. Mean ± s.e.m.; n = 3 independent exposures. g Basal DDB2 levels (normalized to actin) plotted against cisplatin IC_₅₀_ in HT29, Rko, SW480 and HCT116 cells, showing no correlation with cisplatin sensitivity. h Basal DDB2 levels plotted against oxaliplatin IC_₅₀_ in the same cell lines, showing that higher DDB2 levels correspond to higher oxaliplatin IC_₅₀_. For g and h, cells were treated with drug for 2 h and viability (CellTiter-Glo) assessed at 72 h; n = 10. i Dose-response curves of percent survival following oxaliplatin treatment in HCT116, SW480, Rko and HT29 cells by clonogenic assay. Mean ± s.e.m.; n = 6 from two independent experiments. j Percent survival following cisplatin treatment in DDB2-WT versus DDB2-KO U2OS cells. Mean ± s.e.m.; n = 6 from two independent experiments; n.s., not significant by two-way ANOVA at each dose. k Percent survival following oxaliplatin treatment in DDB2-WT versus DDB2-KO U2OS cells, showing significantly reduced survival in DDB2-KO cells with increasing dose. Mean ± s.e.m.; n = 6 from two independent experiments; **P < 0.01, ***P < 0.001, ****P < 0.0001, two-way ANOVA at each dose. l Kaplan–Meier overall survival curves for all TCGA-COAD colorectal cancer patients treated with oxaliplatin, stratified by tumor DDB2 expression. n = 72 per group; p = 0.017, log-rank test. m As in l, for stage IV patients treated with oxaliplatin only. n = 22 per group; p = 0.045, log-rank test.

We next asked whether basal DDB2 protein levels were associated with sensitivity to cisplatin and oxaliplatin, using two complementary assays. First, we used a short-term CellTiter-Glo (CTG) assay to derive the IC_50_ values for each CRC line (Supplementary Fig. S1 a - h). Basal DDB2 levels showed no significant correlation with cisplatin IC_50_ (Fig 1 g; Supplementary Fig. S1 i; R^2^ = 0.039, p = 0.802). In contrast, DDB2 levels correlated positively with oxaliplatin IC_50_, such that DDB2-high lines HCT116 and SW480 were more resistant than DDB2-low lines HT29 and RKO (Fig 1 h; Supplementary Fig. S1 j; R^2^ = 0.977, p = 0.0112). Second, we assessed oxaliplatin sensitivity using a long term clonogenic assay. HCT116 and SW480 cells survived oxaliplatin better across the dose range than HT29 and RKO cells across most of the dose range (Fig. 1 i). The oxaliplatin IC_50_ values derived from this assay again correlated with DDB2 levels (Supplementary Fig. S1 k; R^2^ = 0.915, p = 0.0434). Together, these results suggested that DDB2 protein levels, rather than a wholesale defect in NER capacity, track specifically with oxaliplatin resistance.

Since higher DDB2 protein levels are associated with reduced oxaliplatin sensitivity, we next asked whether DDB2-knockout (KO) cells are more sensitive to oxaliplatin than their wild-type (WT) counterparts. To test this, we treated DDB2-WT and DDB2-KO U2OS cells (Supplementary Fig. S1 l) with cisplatin and oxaliplatin, and compared survival using a clonogenic assay (Fig. 1 j, k). DDB2-KO cells showed no significant difference in survival relative to DDB2-WT cells across a range of cisplatin doses (Fig. 1 j), indicating that loss of DDB2 does not sensitize cells to cisplatin. In sharp contrast, DDB2-KO cells were significantly more sensitive to oxaliplatin than DDB2-WT cells, with the magnitude of this difference increasing at higher drug concentrations (Fig. 1k). We observed IC_50_ values of 5.64 µM and 11.03 µM for cisplatin and oxaliplatin, respectively, for DDB2-WT cells (Supplementary Fig. S1 m). For DDB2-KO cells, we observed IC_50_ values of 3.37 µM and 4.14 µM for cisplatin and oxaliplatin, respectively (Supplementary Fig. S1 m). These results indicate that loss of DDB2 preferentially sensitizes cells to oxaliplatin, in line with the drug-specific association observed in the CRC lines. Finally, to assess whether this relationship extends to clinical outcomes, we analyzed DDB2 expression and survival data from colorectal cancer patients treated with oxaliplatin using the TCGA-COAD cohort. Patients with high tumor DDB2 expression showed significantly worse overall survival than those with low DDB2 expression across the full cohort (n = 72 per group; p = 0.017, log-rank test) (Fig. 1l). This association was also observed within the subset of patients with stage IV disease (n = 22 per group; p = 0.045, log-rank test) (Fig. 1m), suggesting that elevated DDB2 expression is associated with poorer clinical response to oxaliplatin-based therapy even in advanced disease. We also examined other DNA repair proteins; LIG1, LIG3, XPA, XPC and ERCC1, in the same cohort (Supplementary Fig. S1 n, o). Only LIG3 approached statistical significance (p = 0.07, log-rank test) (Supplementary Fig. S1 n), suggesting that the prognostic association is strongest for DDB2 rather than a uniform feature of NER gene expression in CRC.

### DDB2 preferentially recognizes GG oxaliplatin adducts and exhibits higher affinity for oxaliplatin than cisplatin adducts

To determine whether DDB2 directly recognizes cisplatin- and oxaliplatin-induced DNA adducts independent of differences in how these lesions are processed in cells, we used our previously developed single-molecule analysis of DNA-binding proteins from nuclear extracts (SMADNE) assay using the LUMICKS C-trap technology ^28–30^. We first examined whether fluorescently-tagged DDB2 can bind to both cisplatin or oxaliplatin adducts with similar affinities. In this experiment, λ-DNA was treated with oxaliplatin (25 µM) or cisplatin (3.75 µM) for 2 hours to induce approximately the same number and kinds of adducts ^24, 31^. Halo-DDB2 remained bound approximately twice as long on oxaliplatin-treated DNA (1.2 ± 0.03 s) compared with cisplatin-treated DNA (0.6 ± 0.01 s) (Supplementary Fig. S2c, f, g). By measuring how often DDB2 visited a binding site, we obtained an apparent on rate; using the protein concentration in the flow cell, determined by fluorescence with known standards, we calculated true on rates. Obtaining an equilibrium dissociation constant, *K*_D_, by dividing the off rate by the on rate, DDB2 showed approximately 2.7-fold higher affinity for oxaliplatin-treated λ-DNA, with *K*_D_ values of 7.2 ± 0.2 nM and 19.2 ± 0.4 nM for oxaliplatin and cisplatin, respectively. These measurements further support our observation that DDB2 recognizes DNA adducts generated by both platinum drugs, but binds more strongly to oxaliplatin-damaged DNA under the conditions tested.

One drawback to these experiments is that platinum compounds make a spectrum of adducts including: GG, AG, GXG intrastrand adducts and GG interstrand adducts. Also, the sequence context of these adducts could affect DDB2 binding affinities. To overcome these limitations, we built 12 kb DNA fragments containing one oxaliplatin or cisplatin 1,2-intrastrand d(GpG), GG, or one, 1,3-intrastrand d(GpTpG), GTG adduct in the same DNA sequence context directly in the center of the fragment. As with the platinum-treated lambda DNA, we observed that DDB2 recognizes site-specific adducts generated by both platinum drugs. For oxaliplatin, DDB2 bound the GG adduct with a *K*_D_ of 0.46 ± 0.03 nM compared with 2.01 ± 0.03 nM for GTG, representing approximately 4.4-fold higher affinity for the GG adduct than the GTG adduct (Fig. 2c,d and Table 1). In contrast, DDB2 showed approximately 1.8-fold higher affinity for GTG than GG cisplatin adducts, with *K*_D_ values of 11.8 ± 0.4 nM and 21.3 ± 1.44 nM, respectively (Fig. 2g,h and Table 1).

**Fig. 2.**
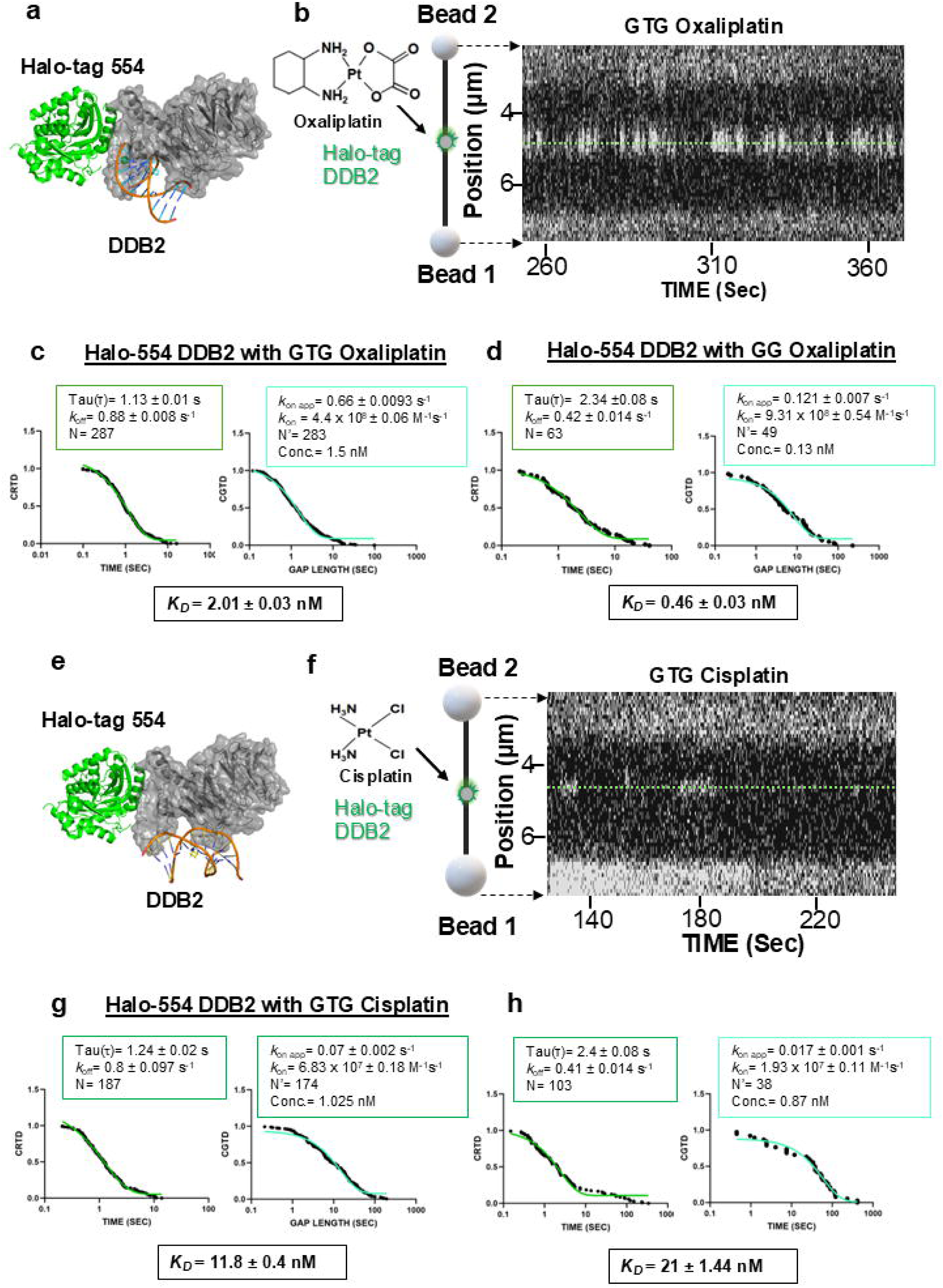
Single-molecule analysis of Halo-tag 554-DDB2 binding to oxaliplatin- and cisplatin-damaged DNA adducts. **a** Representative structural model of N-terminal Halo-tag 554-DDB2 bound to oxaliplatin-damaged DNA. Halo-tag 554 is shown in green and DDB2 in gray. **b** Schematic of the C-trap assay and representative kymograph showing Halo-tag 554-DDB2 binding to a GTG oxaliplatin lesion on DNA tethered between two optically trapped beads. The dashed line shows the DNA path. **c, d** Cumulative residence time (CRTD) and cumulative gap time (CGTD) distributions for DDB2 binding to GTG (c) and GG (d) oxaliplatin adducts. Single-exponential fits are shown in green, and the corresponding binding lifetime, kinetic parameters, and *K*_D_ values are indicated in each panel. **e** Representative structural model of Halo-tag 554 DDB2 bound to cisplatin-damaged DNA. **f** Schematic of the C-trap assay and representative kymograph showing DDB2 binding to a GTG cisplatin lesion. **g, h** CRTD and CGTD analyses of DDB2 binding to GTG (g) and GG (h) cisplatin adducts, respectively. The corresponding kinetic parameters and *K*_D_ values are indicated in each panel.

**Table 1.**
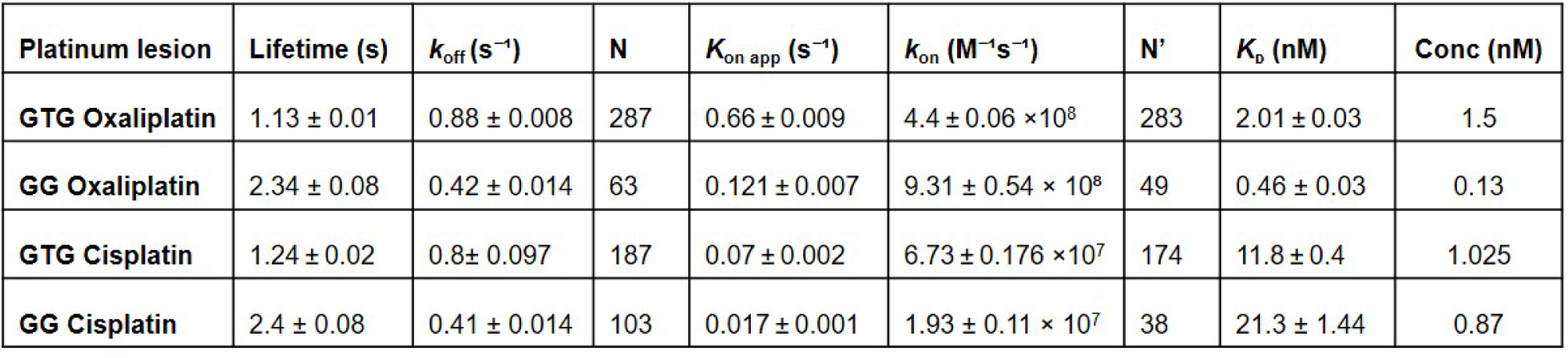
Combined table of binding affinities of Halo-DDB2 to single platinum DNA substrates.

When we compared the two platinum drugs, DDB2 showed approximately 5.9-fold higher affinity for GTG oxaliplatin than for GTG cisplatin adducts. This difference was even greater for GG adducts, where DDB2 exhibited approximately 46-fold higher affinity for oxaliplatin (Fig. 2c,d,g,h). Interestingly, the binding lifetimes were similar for the corresponding adducts. DDB2 remained bound for 1.13 ± 0.01 s on GTG oxaliplatin and 1.24 ± 0.02 s on GTG cisplatin, while the lifetimes for GG oxaliplatin and cisplatin were 2.34 ± 0.08 s and 2.4 ± 0.08 s, respectively. Thus, the differences in affinity appear to arise largely from differences in the association kinetics of these lesions in this sequence context rather than how long DDB2 remains bound. Among the four adducts examined, DDB2 had the highest affinity for GG oxaliplatin adducts.

### Oxaliplatin, but not cisplatin treatment, drives DDB2 accumulation in the nucleolus

Oxaliplatin has been reported to kill cells largely by inducing ribosome biogenesis and nucleolar stress rather than through the canonical DNA damage response that underlies cisplatin toxicity ^32–35^, a mechanistic distinction that could explain why loss of DDB2 sensitizes cells to oxaliplatin but not cisplatin (Fig. 1j, k). Having established that DDB2 binds oxaliplatin and cisplatin DNA adducts in vitro (Fig. 2), we reasoned that the drug-specific phenotype of DDB2-deficient cells is more likely explained by a difference in how DDB2 behaves inside the cell after each treatment, rather than by a difference in lesion recognition. Because nucleolar stress is a hallmark of the oxaliplatin response ^26, 34, 36^, we hypothesized that DDB2 itself relocalizes to the nucleolus following oxaliplatin, but not cisplatin, treatment.

To test this hypothesis, we transiently transfected the U2OS cells stably expressing mNeonGreen-tagged DDB2 ^3^ with the nucleolar marker protein pHcRed-nucleophosmin 1 (NPM1), then imaged cells by confocal microscopy at a single fixed time point or by live time-lapse imaging over 6 h following drug addition (Fig. 3 and 4). To eliminate dosage or potency effects, both cisplatin and oxaliplatin were administered at their IC_₅₀_ dosage obtained from clonogenic assay (5.64 µM for cisplatin and 11.03 µM oxaliplatin; Fig 1j,k). In untreated cells and in cells treated with cisplatin for 6 h, mNeonGreen-DDB2 remained diffusely distributed throughout the nucleus and was excluded from NPM1-marked nucleoli (Fig. 3b). In marked contrast, oxaliplatin treatment caused DDB2 to redistribute into a discrete focus that co-localized with NPM1, indicating that DDB2 specifically accumulates within the nucleolus in response to oxaliplatin (Fig. 3b). Quantification of nucleolar DDB2 intensity across the cell population confirmed this: oxaliplatin-treated cells showed a 2.1-fold-increase in mean nucleolar DDB2 signal relative to untreated controls (P=0.0005, unpaired Student’s t-test), comparable to the increase produced by Actinomycin D, a RNA Pol I inhibitor and a well-characterized inducer of nucleolar stress ^37, 38^, whereas cisplatin-treated cells were indistinguishable from untreated controls (Fig. 3c).

**Fig 3.**
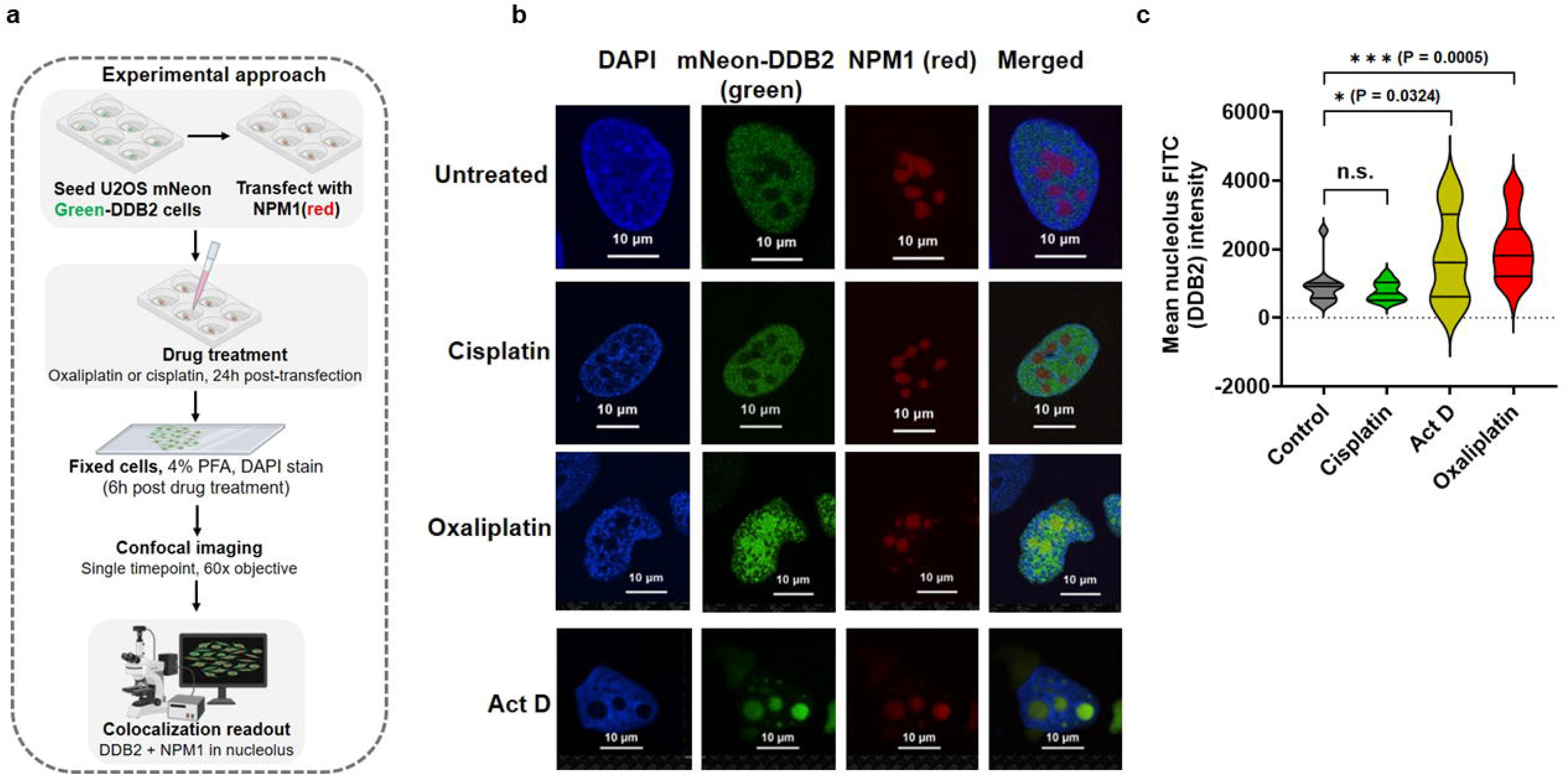
Oxaliplatin, but not cisplatin treatment, induces DDB2 accumulation in the nucleolus with concomitant NPM1 relocalization. **a** Schematic of the experimental workflow. U2OS cells stably expressing mNeonGreen-tagged DDB2 (mNeonGreen-DDB2) were transiently transfected with NPM1-pHcRed and, 24 h later, treated with oxaliplatin or cisplatin. Cells were fixed at 6 h post-treatment. The illustrations were created in BioRender by Kumar, V. (2026). https://BioRender.com/vfmf6ow. **B** Representative confocal images of fixed cells showing DAPI (nuclear DNA, blue), mNeonGreen-DDB2 (green), NPM1-pHcRed (red), and merged channels in untreated cells and cells treated with cisplatin or oxaliplatin for 6 h. In untreated and cisplatin-treated cells, DDB2 remains diffusely nuclear while NPM1 marks discrete nucleoli. Oxaliplatin treatment induces DDB2 redistribution into a single NPM1-positive nucleolar focus, with DDB2 and NPM1 co-localizing in the merged image, indicative of nucleolar stress. Act D: Actinomycin D. Scale bars, 10 µm. **c** Quantification of mean nucleolar DDB2 (FITC) fluorescence intensity per cell in untreated, cisplatin-treated, Actinomycin D (Act D; positive control for nucleolar stress) treated, and oxaliplatin-treated cells. Data are shown as violin plots with median and quartiles. Number of nucleoli analyzed: n = 72 for untreated; n = 68 for cisplatin; n = 228 for oxaliplatin; n = 111 for Act D group. Number of nuclei analyzed: n = 12 for untreated; n = 16 for cisplatin; n = 38 for oxaliplatin; n = 30 for Act D group. NS = not significant, *P < 0.05, and ***P < 0.001 by unpaired t-test.

**Fig 4.**
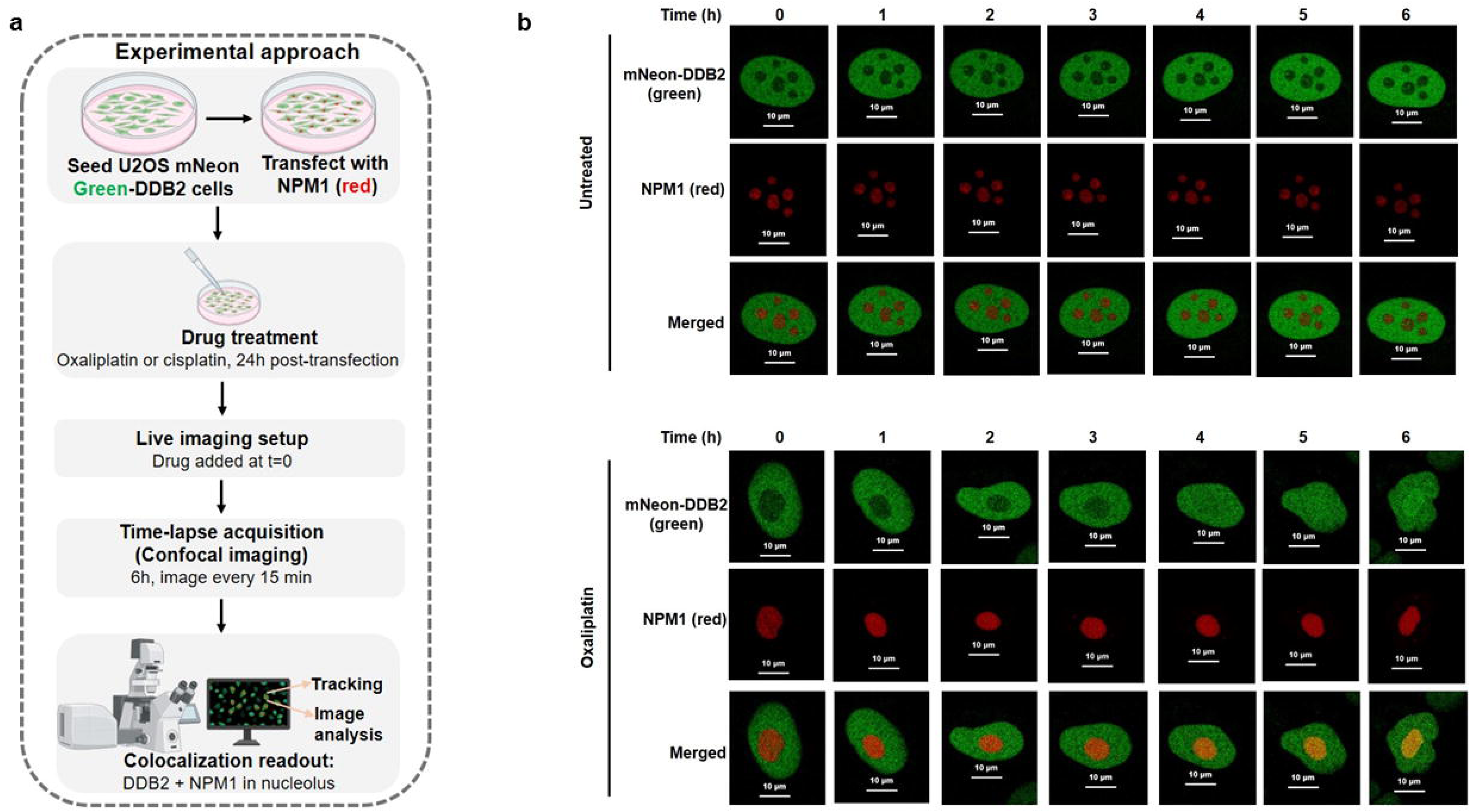
Time-lapse live-cell imaging of mNeonGreen-DDB2 (green) cell, transiently transfected with NPM1-pHcRed (red). **a** Schematic of the experimental workflow. U2OS cells stably expressing mNeonGreen-tagged DDB2 (mNeonGreen-DDB2) were transiently transfected with NPM1-pHcRed and, 24 h later, treated with oxaliplatin or cisplatin, and imaged live by time-lapse confocal microscopy every 15 min for 6 h beginning at drug addition. The illustrations were created in BioRender by Kumar, V. (2026). https://BioRender.com/1n3582a. **b** Live-cell imaging of mNeonGreen-DDB2 (green) and NPM1-pHcRed (red) in untreated and oxaliplatin-treated cells, imaged every 15 min for 6 h (representative frames shown at 1 h intervals). In untreated cells, DDB2 remains diffusely nuclear and excluded from NPM1-marked nucleoli throughout the time course. Following oxaliplatin treatment, DDB2 progressively accumulates within the NPM1-positive nucleolus, with clear co-localization evident by 4 h and persisting through 6 h (merged panels), demonstrating that nucleolar recruitment of DDB2 is a dynamic, time-dependent response to oxaliplatin. Corresponding time-lapse movies are provided in Supplementary videos 1 and 3.

We next asked whether nucleolar accumulation of DDB2 develops progressively over time (Fig. 4a). Live cell time-lapse imaging showed that in untreated and cisplatin-treated cells, DDB2 remained nuclear and excluded from NPM1-positive nucleoli throughout the 6 h imaging window (Fig. 4b, Supplementary Fig. S3 and Supplementary video 1 and 2). Following oxaliplatin addition, however, DDB2 progressively accumulated within the NPM1-marked nucleolus over the course of imaging, with clear co-localization evident by 4 h and persisting through 6 h (Fig. 4b and Supplementary video 3). Together, these data show that DDB2 undergoes a drug-specific, time-dependent relocalization to the nucleolus in response to oxaliplatin, positioning DDB2 at the site of oxaliplatin-induced nucleolar stress rather than solely at sites of direct DNA lesion recognition.

### DDB2 repair activity, not the DDB2 protein itself, is required to protect cells from oxaliplatin

Having established that DDB2 accumulates in the nucleolus specifically after oxaliplatin, but not cisplatin, within 4-6 hr of treatment (Fig. 3 and 4), we determined whether DDB2’s entry into the nucleolus provides a protective cell survival effect by its repair activity or relieving nucleolar stress through another mechanism (Fig. 5a). To distinguish between these possibilities, we compared WT DDB2 to a DDB2 K244E variant. This amino acid substitution on the DNA-binding surface of DDB2 causes the skin cancer-prone syndrome xeroderma pigmentosum complementation group E (XP-E) in patients ^12^ and renders DDB2 defective for damage recognition ^39^ (Fig. 5b). We reconstituted DDB2-KO U2OS cells with either DDB2-WT or DDB2-K244E, together with NPM1 (Supplementary Fig. 4a), reasoning that if DDB2’s protective role against oxaliplatin were independent of its repair activity, DDB2-K244E should rescue cells similarly to DDB2-WT; conversely, if repair activity is required, DDB2-K244E-expressing cells should instead behave like DDB2-KO cells (Fig. 5a).

**Fig. 5.**
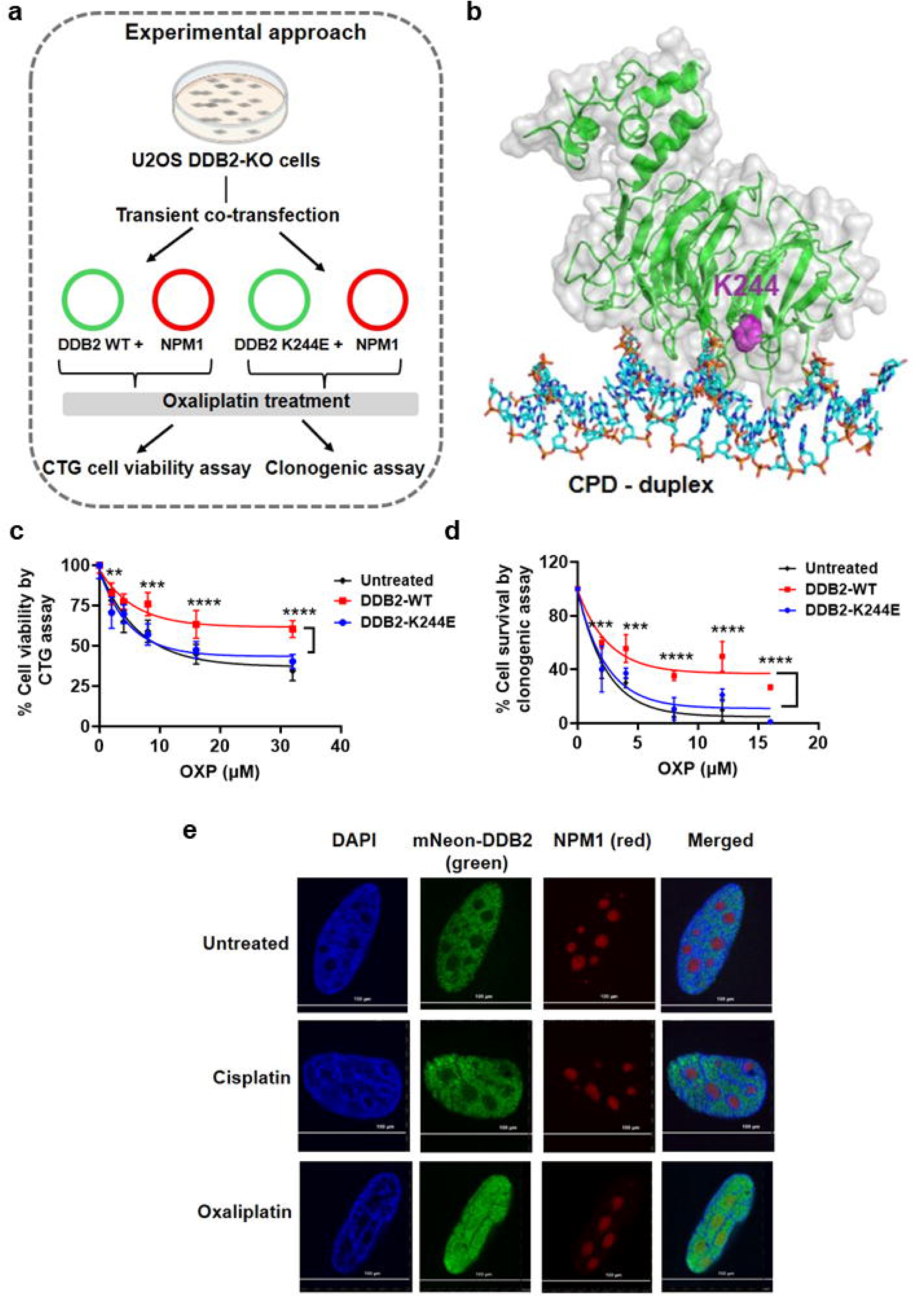
DDB2 repair function is important for DDB2-induced oxaliplatin response. **a** Schematic of the experimental approach. U2OS DDB2-knockout (DDB2-KO) cells were transiently co-transfected with plasmids encoding NPM1 together with either wild-type DDB2 (DDB2-WT) or the repair-deficient DDB2 K244E variant, followed by oxaliplatin treatment. The illustrations were created in BioRender by Kumar, V. (2026). https://BioRender.com/zm917va. **b** Structural model of DDB2 showing binding to cyclobutane pyrimidine dimers (CPD) and the position of the K244 residue (in magenta); the K244E mutation causes the skin cancer-prone syndrome xeroderma pigmentosum complementation group E (XP-E). The structure was retrieved from PDB Id:4A08. **c** Clonogenic survival of DDB2-KO cells expressing DDB2-WT or DDB2-K244E, treated with increasing concentrations of oxaliplatin for 2 h, relative to untreated controls. Data are presented as mean ± s.e.m. from n = 6 from 2 biologically independent experiments. **P < 0.001, ****P < 0.0001 by two-way ANOVA with Sidak’s multiple comparisons test, DDB2-WT vs. DDB2-K244E. **d** Cell viability (CellTiter-Glo; CTG) of the same cell populations following oxaliplatin treatment, shown as % viability relative to untreated controls. The cells were treated with oxaliplatin for 2 h, and CTG readings were taken after 48 h. Data are mean ± s.e.m. from n = 8, from two independent experiments. **P < 0.01, ***P < 0.001, ****P < 0.0001 by two-way ANOVA with Sidak’s multiple comparisons test, DDB2-WT vs. DDB2-K244E. **e** Representative confocal images of DDB2-KO cells co-expressing mNeonGreen-DDB2 K244E and NPM1-pHcRed, showing DAPI (nuclear DNA, blue), mNeon-DDB2 (green), NPM1 (red), and merged channels in untreated cells and cells treated with cisplatin or oxaliplatin for 6 h. As with DDB2-WT (Fig. 3b), oxaliplatin but not cisplatin induces redistribution of the K244E variant into the NPM1-positive nucleolus. Scale bars, 100 µm.

Consistent with a repair-dependent mechanism, the CTG cell viability assays showed that reconstitution with DDB2-WT substantially rescued the viability of DDB2-KO cells across a range of oxaliplatin concentrations, whereas cells expressing DDB2-K244E showed no meaningful improvement in cell viability relative to untreated DDB2-KO controls (Fig. 5c). This result was corroborated using an orthogonal, clonogenic survival assays in which DDB2-WT-expressing cells maintained significantly higher cell survival than DDB2-K244E-expressing cells following oxaliplatin treatment across all tested doses (Fig. 5d and Supplementary Fig. 4b).

Finally, we asked whether the K244E mutation also disrupts the oxaliplatin-induced nucleolar recruitment of DDB2 we observed previously (Fig. 3 and 4), or whether nucleolar localization and repair function are mechanistically separable. Confocal imaging showed that, similar to DDB2-WT, the DDB2-K244E variant still relocalized into the NPM1-positive nucleolus following oxaliplatin, but not cisplatin, treatment (Fig. 5e). Together, these data indicate that the repair-competent DNA-binding activity of DDB2, rather than the protein’s mere presence in the nucleolus, is required to protect cells from oxaliplatin.

### Pharmacological degradation of DDB2 sensitizes CRC cells to oxaliplatin

After establishing that DDB2 protein levels track with oxaliplatin resistance or response (Fig. 1) and that DDB2’s repair activity, rather than nucleolar localization alone, is required to protect cells from oxaliplatin (Fig. 5), we next asked whether pharmacologically altering DDB2 activity could resensitize resistant CRC cells to oxaliplatin. DDB2 is a substrate receptor for the CRL4^DDB2^ E3 ubiquitin ligase, whose activity is governed by a cycle of neddylation, mediated by the NEDD8-activating enzyme (NAE), and deneddylation, mediated by the COP9 signalosome (CSN) (Fig. 6a). We reasoned that disrupting either arm of this cycle would sensitize cells to oxaliplatin. We tested this hypothesis using either the NAE inhibitor pevonedistat (NAEi/MLN4924), which blocks UV-DDB activation, or the CSN5 inhibitor CSN5i-3, which drives CRL4^DDB2^ into a hyperactive state, triggering autoubiquitination and destruction of DDB2 itself ^3^. Treatment of U2OS cells, which expresses high basal DDB2 (Fig. 1), with CSN5i-3 produced a time- and dose-dependent loss of DDB2 protein, accompanied by induction of the CDK inhibitor p21 (Fig. 6b). We then tested whether DDB2 degradation could sensitize HCT116 CRC cells to oxaliplatin. HCT116 cells express the highest DDB2 protein levels and are least sensitive to oxaliplatin (Fig. 1). In cell viability assays, both NAEi and CSN5i-3 modestly reduced viability as single agents (reduced cell viability by ∼20% at highest doses), but each dramatically enhanced oxaliplatin cytotoxicity in combination, shifting the dose-response curves substantially leftward relative to oxaliplatin alone (Fig. 6c, d). We used the Chou-Talalay method ^40, 41^ to determine synergy in HCT116 cells. The data were analyzed with Compusyn software (freely downloadable from https://www.combosyn.com/) to generate the Fa-CI (Fa: factor affected; CI: combination index) curves. We observed CIs of 0.0335, 0.0387 and 0.0392 for C1, C2, and C3 combinations at 72 h for oxaliplatin + NAEi. We observed CIs of 0.3113, 0.2461, and 0.3374 for C1, C2, and C3 combinations at 72 h for oxaliplatin + CSN5i-3. These interactions, using the Chou-Talalay combination index method, confirmed that oxaliplatin combined with either NAEi or CSN5i-3 produced strongly synergistic effects (CI < 1) across nearly the entire range of fraction affected (Fig. 6c, d). This synergy was further supported by using an orthogonal, longer-term clonogenic survival assay: combined treatment with oxaliplatin and either DDB2 inhibitor (NAEi : DDB2 inactivator and CSN5i-3: DDB2 degrader) ^3^ nearly abolished colony formation at doses that had only a modest effect as single agents (Fig. 6e and Supplementary Fig. S5). Together, these results demonstrate that pharmacological degradation of DDB2 via CRL4^DDB2^ hyperactivation ^3^ is sufficient to overcome DDB2-mediated oxaliplatin resistance, providing proof-of-principle for a biomarker-driven combination strategy in CRC.

**Fig. 6.**
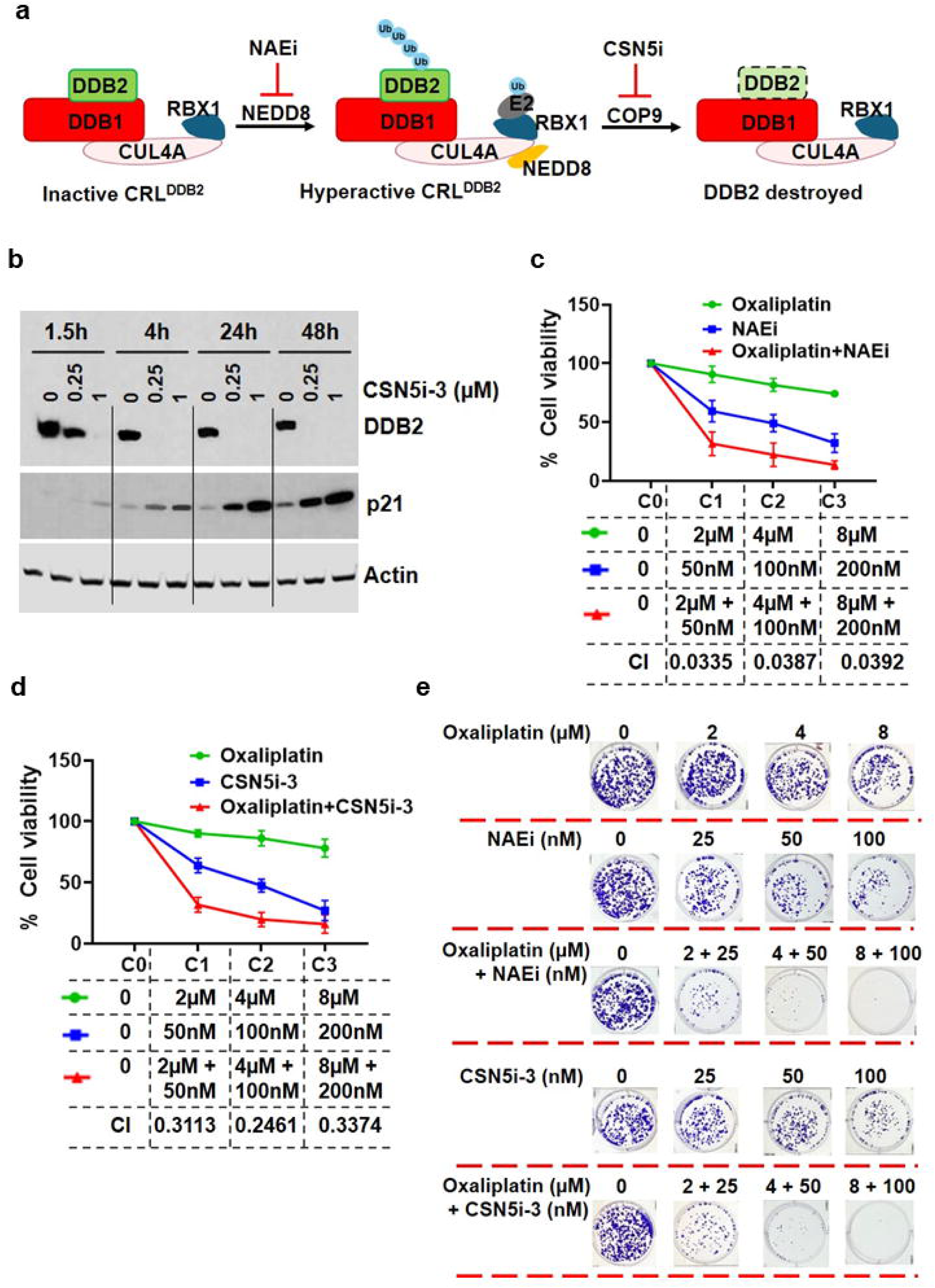
Pharmacological inhibition of DDB2 activity synergizes with oxaliplatin in colorectal cancer cells. **a**, Schematic of the CUL4A-RBX1-DDB1-DDB2 E3 ubiquitin ligase complex (CRL^DDB2^) and its cycle of activation and inactivation through NEDD8 conjugation and removal. In the basal state, the inactive CRL^DDB2^ complex (CUL4A, RBX1, DDB1, and the substrate receptor DDB2) lacks NEDD8 modification. Conjugation of NEDD8 onto CUL4A, catalyzed by the NEDD8-activating enzyme (NAE) together with an E2 enzyme, converts the complex into its hyperactive, neddylated form (Hyperactive CRL^DDB2^), which promotes ubiquitination of substrates including DDB2 itself. This activating step is blocked by the NAE inhibitor NAEi. Continued ligase activity in the neddylated state leads to DDB2 ubiquitination and proteasomal degradation (DDB2 destroyed; dashed outline indicates loss of DDB2). Deneddylation of CUL4A by the COP9 signalosome (CSN), via its catalytic subunit CSN5, returns the complex to the inactive state, completing the cycle; this step is blocked by the CSN5 inhibitor CSN5i. Ub, ubiquitin. The illustrations were created in BioRender by Kumar, V. (2026). https://BioRender.com/yhok1h8. **b** Western blot of DDB2 and p21 in U2OS cells treated with CSN5i-3 (0, 0.25 or 1 µM) for 1.5, 4, 24 or 48 h. β-actin, loading control. **c**, Viability of HCT116 cells treated with oxaliplatin, NAEi, or oxaliplatin + NAEi at the indicated paired concentrations (C0-C3), for 2 h for oxaliplatin and 72 h for NAEi or CSN5i-3, measured by CellTiter-Glo. Data are mean ± s.e.m., n = 12 from 3 independent experiments. **d**, As in **c**, for oxaliplatin, CSN5i-3, or oxaliplatin + CSN5i-3. The combination index (CI) was calculated by the Chou-Talalay method. CI < 1, synergistic; CI = 1, additive; CI > 1, antagonistic. **e**, Representative clonogenic assay images of HCT116 cells treated with oxaliplatin, NAEi, or CSN5i-3 alone, or the indicated combinations, stained with crystal violet. n = 3 (In the clonogenic assay, for the full triplicate plate picture, the reader is referred to Supplementary Fig. 6 e; only a cropped version is presented here).

### Elevated DDB2 expression predicts oxaliplatin/FOLFOX resistance and independently prognosticates clinical outcome

To determine whether the relationship between DDB2 and oxaliplatin resistance established in our cell-based and TCGA analyses (Fig. 1) extends to an independent clinical cohort, we examined *DDB2* mRNA expression in a combined cohort of 86 CRC patients treated with first-line FOLFOX/oxaliplatin (GEO datasets GSE83129, GSE28702 and GSE69657)^42–44^, stratified by response as Responders (n = 45) or Non-responders (n = 41). Non-responders showed nearly two-fold higher normalized *DDB2* transcript levels than responders, a difference that was highly significant (Fig. 7a; p < 0.0001, Welch’s *t*-test). To determine whether this association was independent of other clinical variables, we built a multivariate Cox proportional hazards model for progression-free survival incorporating *DDB2* expression, age at diagnosis, pathological stage, and microsatellite status. *DDB2* expression remained a strong, independent predictor of shorter progression-free survival (hazard ratio (HR) = 2.14 per log_₂_ increase, 95% CI (confidence interval) 1.58-2.91, P < 0.001), comparable in magnitude to pathological stage (HR = 1.85 for stage IV vs. III, p = 0.016), whereas age and microsatellite status were not independently significant (Fig. 7b). These results indicate that tumor *DDB2* levels carry independent prognostic value for oxaliplatin-based therapy outcomes, beyond established clinicopathological risk factors.

**Fig. 7.**
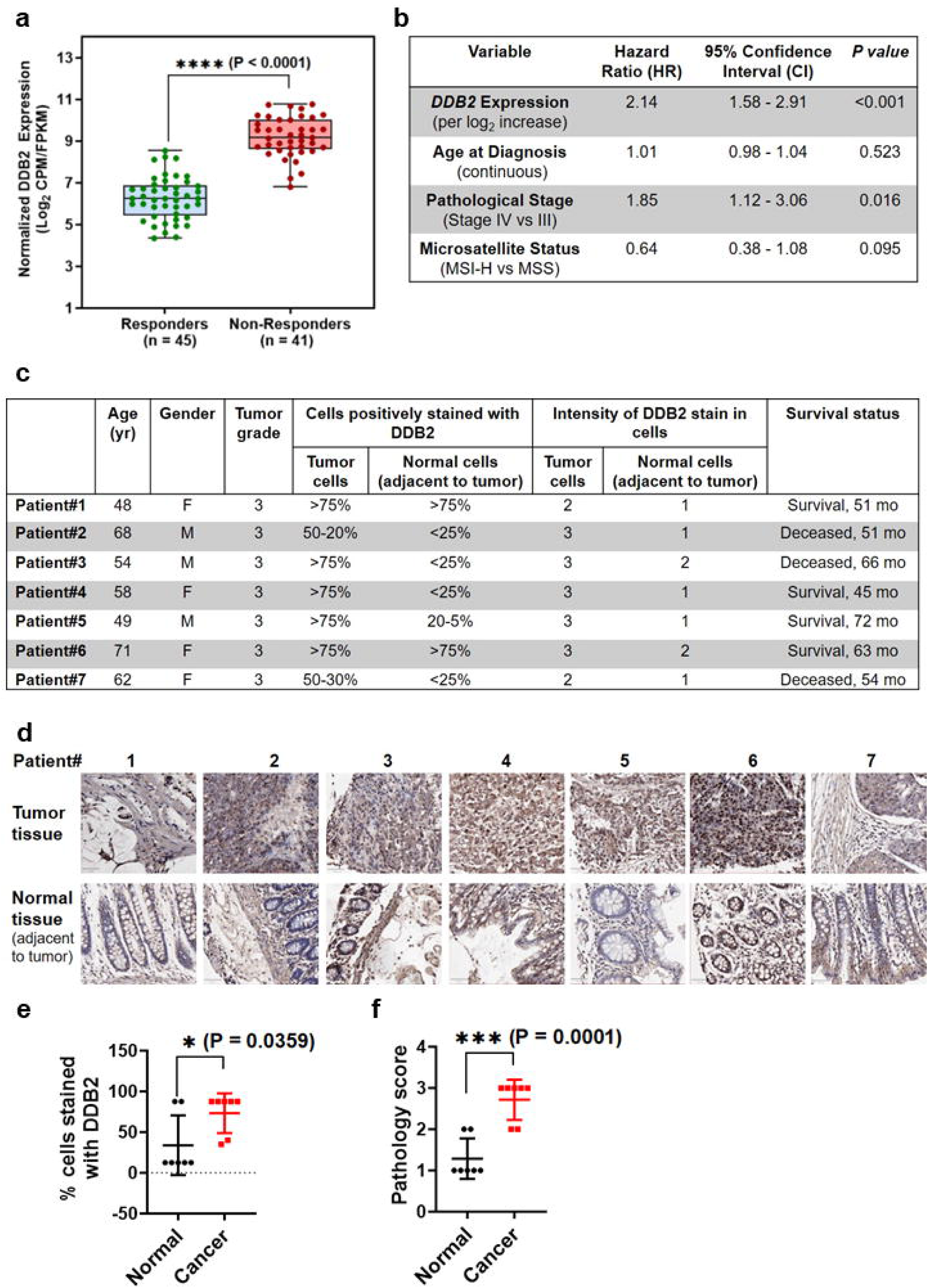
Elevated *DDB2* expression is associated with oxaliplatin/FOLFOX resistance and independently predicts poor prognosis in colorectal cancer patients. **a** Normalized *DDB2* mRNA expression (log_₂_ CPM/FPKM) in tumors from patients with advanced/metastatic colorectal cancer (CRC) treated with first-line FOLFOX/oxaliplatin. Patients documented as achieving a Complete Response (CR) or Partial Response (PR) were classified as Responders (n=45). Patients exhibiting Stable Disease (SD) or Progressive Disease (PD) were classified as Non-Responders (n=41). GEO datasets used: GSE83129, GSE28702, GSE69657. \*\*\**P* < 0.0001, two-tailed unpaired Student’s *t*-test with Welch’s correction. CPM: Counts Per Million; FPKM: Fragments Per Kilobase of transcript per Million mapped reads. b Multivariate Cox proportional hazards regression for progression-free survival in the cohort from a, adjusted for *DDB2* expression (per log_₂_ increase), age at diagnosis, pathological stage (IV vs. III), and microsatellite status (MSI-H vs. MSS). Hazard ratios (HR), 95% confidence intervals (CI), and *P* values are shown for each covariate. **c** Clinicopathological characteristics and semi-quantitative DDB2 immunohistochemistry (IHC) scoring in a paired tissue microarray (TMA) of seven grade 3 colon adenocarcinomas (tumor and matched adjacent normal tissue). The percentage of DDB2-positive cells and staining intensity (0, none; 1, weak; 2, moderate; 3, strong) were scored for tumor and adjacent normal tissue per patient, with corresponding survival status/follow-up (months). **d** Representative DDB2 IHC images (20× magnification) of tumor and matched adjacent normal tissue from the seven patients in **c**. **e** Quantification of the percentage of DDB2-positive cells in normal versus paired tumor tissue from **c**, **d** (n = 7 patient pairs). Cells scored within a range (<25% or >75%) were assigned the midpoint of that range for statistical analysis. *p = 0.0359, two-tailed unpaired Student’s *t*-test. **f** Quantification of DDB2 staining intensity (pathology score) in normal versus paired tumor tissue from **c**, **d** (n = 7 patient pairs). ***p = 0.0001, two-tailed unpaired Student’s *t*-test. Data in **e**, **f** are presented as mean ± s.e.m.

Since mRNA and protein levels do not always correlate, and IHC-based markers are more readily translatable to clinical practice, we next asked whether DDB2 protein levels were similarly elevated in colon tumor tissue. We performed DDB2 immunohistochemistry on a paired tissue microarray comprising seven grade 3 colon adenocarcinomas with matched adjacent normal tissue (Fig. 7c, d and Supplementary Fig. S6). Semi-quantitative scoring showed that both the percentage of DDB2-positive cells (Fig. 7e) and DDB2 staining intensity (Fig. 7f) were significantly higher in tumor than in matched normal tissue (p = 0.0359 and p = 0.0001, respectively, unpaired Student’s *t*-test). Together, these clinical and histological analyses demonstrate that elevated DDB2 expression, at both the transcript and protein level, is associated with reduced oxaliplatin/FOLFOX responsiveness and worse prognosis in colorectal cancer, supporting DDB2 as a candidate predictive biomarker for oxaliplatin-based therapy.

## Discussion

Colorectal cancer (CRC) is the second most common cause of cancer death in the United States, and colon cancer is the most common type of gastrointestinal cancer. According to the American Cancer Society’s (ACS) estimates, in people younger than 50 years of age, rates have increased by 3% per year from 2013 to 2022. Oxaliplatin, as part of FOFLOX treatment, is used as the standard of care in CRC. In this study, we set out to resolve why loss of DDB2 selectively sensitizes cancer cells to oxaliplatin but not to cisplatin, despite both platinum drugs generating similar major classes of DNA intrastrand crosslinks. Using single-molecule C-trap analysis, we found that DDB2 binds to both oxaliplatin and cisplatin GTG and GG adducts, ruling out a simple lesion recognition defect as the basis for this drug-specific phenotype. Live-cell and fixed-cell imaging instead revealed that oxaliplatin, but not cisplatin, drives progressive and selective accumulation of DDB2 within the NPM1-marked nucleolus, and that this nucleolar recruitment occurs independently of DDB2’s repair activity, since the repair-deficient K244E variant still relocalizes to the nucleolus yet fails to protect cells from oxaliplatin, indicating that DDB2’s DNA binding and repair function, once within the nucleolus, rather than nucleolar localization per se, underlies its protective role. Extending these mechanistic findings to patient-relevant contexts, we showed that basal DDB2 protein levels vary widely across CRC cell lines and track specifically with oxaliplatin, not cisplatin, resistance/response; that elevated tumor DDB2 expression, both at the transcript level in an independent cohort of FOLFOX/oxaliplatin-treated patients and at the protein level by immunohistochemistry, is associated with treatment non-response; and that high tumor DDB2 predicts significantly worse overall survival in the TCGA-COAD cohort, independent of pathological stage. Finally, we found that pharmacologically altering DDB2 activity by NAE inhibitor pevonedistat or the CSN5 inhibitor CSN5i-3 strongly synergized with oxaliplatin to re-sensitize high-DDB2 CRC cells, providing proof-of-principle for a biomarker-driven combination strategy. Together, these findings establish that DDB2 is redirected to, and performs a protective repair function within, the nucleolus specifically in response to oxaliplatin. This raises the possibility that this DDB2-dependent protective mechanism may be exploited therapeutically by pairing oxaliplatin with pharmacological DDB2 degraders in DDB2-high tumors.

Since DDB2 recognizes distortions in DNA, differences in the local DNA conformations induced by oxaliplatin and cisplatin may influence DDB2 binding. Using our single-molecule analysis of DNA-binding proteins from nuclear extracts (SMADNE) assay ^28, 29^ clearly indicated that DDB2 binds oxaliplatin and cisplatin intrastrand crosslinks with high affinity. The binding affinities, *K*_D_, to site-specific DNA adducts followed a strong-to-weak order: GG-oxaliplatin, GTG-oxaliplatin, GTG-cisplatin, and finally GG-cisplatin (Fig. 2 and Supplementary Fig. 2; Table 1). Despite these differences in affinity, the binding lifetimes were similar for corresponding adducts, suggesting that differences in association kinetics contribute to the ehanced recognition of oxaliplatin-induced DNA damage. Early gel-shift studies with purified UV-DDB found strong binding to (6-4) photoproduct and abasic-site duplexes, as well as duplexes containing cisplatin adducts or CPD, and these studies suggested that UV-DDB recognizes a kink, or a readily bendable site, in the DNA ^13, 14^.

Our live-cell and fixed-cell imaging data identified important mechanistic differences in the response to the two platinum compounds: oxaliplatin, but not cisplatin, drives progressive and selective accumulation of DDB2 within the NPM1-marked nucleolus (Fig. 3 and 4). This finding directly links DDB2 migration into the nucleolus to either the oxaliplatin- or Actinomycin-induced nucleolar stress pathway. Oxaliplatin-linked nucleolar stress is proposed to underlie the distinct cytotoxic mechanism of oxaliplatin relative to cisplatin ^26, 27^. Consistent with this, oxaliplatin at ∼10 µM inhibits Pol I transcription within 90 min and relocalizes NPM1 by ∼4 h, before any appreciable γH2AX signal, whereas cisplatin elicits a stronger DNA damage response with little nucleolar disruption ^35^. Nucleolar stress by oxaliplatin depends on the 1R,2R-DACH carrier ligand ^32^, involves ATM/ATR signaling in the absence of detectable nucleolar DNA damage ^27^, and has been linked to partitioning of oxaliplatin into nucleoli, where it modifies fibrillarin and causes nucleolar demixing in HCT116 and U2OS cells ^33^. Our data indicate that oxaliplatin, by provoking nucleolar stress, provides an opportunity for DDB2 to enter into the nucleolus to initiate DNA repair. Genome-wide mapping shows that cisplatin damage formation is determined largely by GG frequency^45^, and human rDNA is GG-rich ^46^, which would make it an abundant target for oxaliplatin-GG adducts. This model reframes DDB2 not simply as a genome-wide damage sensor, but as a context-dependent responder that is mobilized to distinct subnuclear compartments according to the type of genotoxic or organellar stress imposed, providing a cellular explanation for the drug-specific phenotype of DDB2-deficient cells reported previously and recapitulated in this study (Fig. 1j, k).

A key finding distinguishes this nucleolar recruitment from a purely repair-independent “nucleolar guardian” function. The repair-deficient K244E variant of DDB2 ^39^ that causes xeroderma pigmentosum complementation group E (XP-E) in patients^12^ still relocalizes to the nucleolus after oxaliplatin, but does not protect DDB2 KO from oxaliplatin sensitivity (Fig. 5c, d, e). Together, these results indicate that DDB2’s protective function requires its DNA damage recognition capacity once it reaches the nucleolus, not merely its physical presence within the organelle. We propose that DDB2 initiates active global genome nucleotide excision repair (GG-NER) on nucleolar or ribosomal DNA adducts, and that this repair activity, rather than passive sequestration of damage or a repair-independent scaffolding role, mitigates oxaliplatin induced nucleolar stress and preserves cell viability.

Higher levels of the DDB2 protein are linked to poorer treatment outcomes and resistance to the drug oxaliplatin. Cell lines with the highest basal DDB2 protein were the most oxaliplatin resistant (Fig. 1), and in two independent clinical contexts, elevated tumor *DDB2* tracked with worse response: transcript levels were nearly two-fold higher in FOLFOX/oxaliplatin non-responders than responders and remained an independent predictor of shorter progression-free survival after adjustment for stage and microsatellite status (Fig. 7a, b), and DDB2 protein was likewise elevated in tumor relative to matched normal tissue by IHC (Fig. 7e, f), while high tumor DDB2 was associated with significantly worse overall survival in the TCGA-COAD cohort (Fig. 1l, m). We suggest that DDB2’s canonical role as an initiator of global genome repair, tumor-suppressive during carcinogenesis, becomes co-opted once a tumor has formed, so that intact nucleolar surveillance and repair instead confer a survival advantage against oxaliplatin-induced stress. This duality underscores why DDB2 protein or transcript abundance, rather than mutational status, is the biologically and clinically relevant biomarker in the therapeutic setting. The recognition that both the amounts of DDB2 activity and oxaliplatin lesion detection are important determinants of oxaliplatin resistance pointed us toward a therapeutic strategy: rather than targeting DDB2’s adduct binding directly, we inactivated the protein with the NAE inhibitor, pevonedistat ^47^, or degrading DDB2 by hyperactivating the CRL4^DDB2^ ligase through the neddylation cycle with the CSN5 inhibitor, CSN5i-3 ^48^. Treatment with either compound (Fig. 6a) produced strongly synergistic killing with oxaliplatin in high-DDB2 HCT116 cells across viability and clonogenic assays (Fig. 6c-e). Because pevonedistat is already in clinical development, this combination strategy could in principle be advanced relatively quickly. In addition, our data provide proof-of-principle that DDB2 levels can be used both to identify patients likely to be oxaliplatin resistant and to guide a rational, biomarker-informed combination approach to overcoming that resistance. Finally, this synergistic effect on cell killing suggests that, if this combination therapy is used in the clinic, lower doses of oxaliplatin would be effective, thus lowering its toxicity.

Collectively, our findings resolved the apparent contradiction between DDB2’s in vitro lesion binding affinity and its drug-specific role in cellular resistance. Our working model is that DDB2 is redirected to the nucleolus and performs repair on ribosomal genes specifically in response to oxaliplatin. This study further establishes DDB2 as a tractable, biomarker-guided target: elevated tumor DDB2 predicts oxaliplatin/FOLFOX resistance and poor prognosis, and pharmacological degradation of DDB2 restores oxaliplatin sensitivity (Fig. 8 and Supplementary video 4). These results provide a rationale for prospective, biomarker-stratified trials combining oxaliplatin-based regimens with clinically available neddylation pathway inhibitors in DDB2-high colorectal cancer.

**Figure 8.**
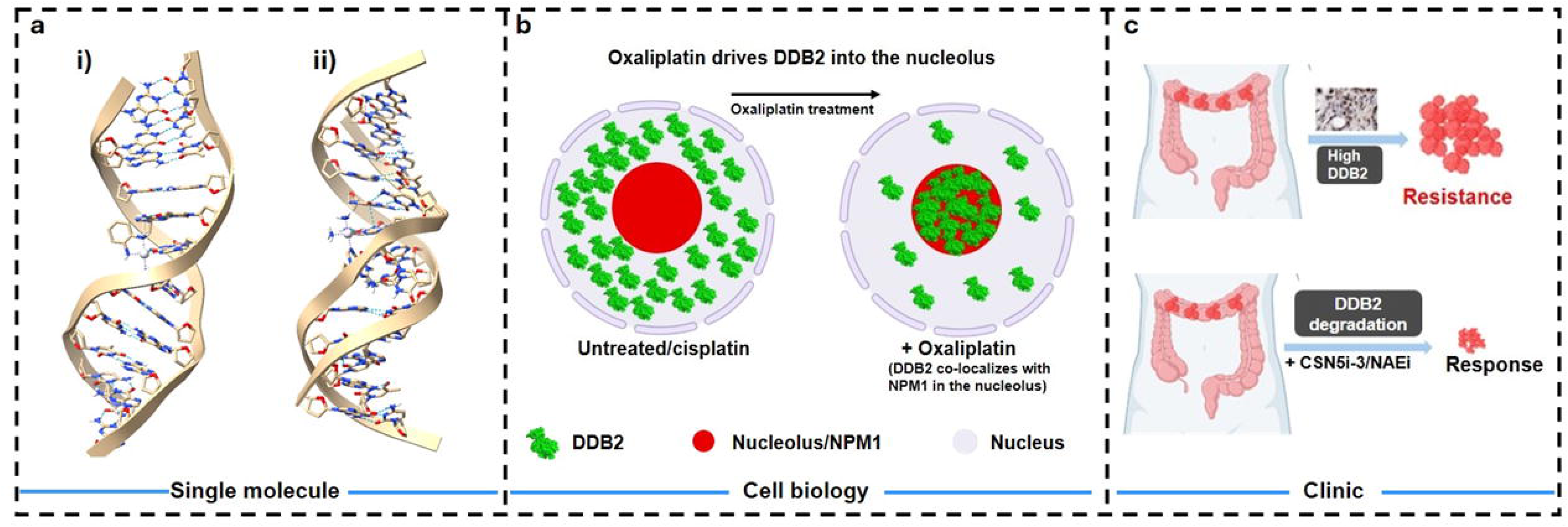
Summary and working model. **a i)** Structure of oxaliplatin GG lesion, PDB ID: 2KOU. **ii)** Structure of cisplatin GTG lesion, PDB ID: 1DA5. **b** Schematic illustrating that, unlike cisplatin, oxaliplatin treatment drives redistribution of DDB2 from the nucleoplasm into the nucleolus, where it co-localizes with the nucleolar marker NPM1. In untreated or cisplatin-treated cells, DDB2 (green) remains dispersed throughout the nucleus; following oxaliplatin treatment, DDB2 accumulates within the nucleolus (red). **c** Proposed model of DDB2-mediated oxaliplatin resistance and its reversal. Elevated DDB2 levels in colorectal cancer cells promote resistance to oxaliplatin. Pharmacological degradation of DDB2 using CSN5i-3 or an NAE1 inhibitor (NAEi) restores sensitivity to oxaliplatin, converting a resistant phenotype into a therapeutic response. The illustrations were created in BioRender by Kumar, V. (2026). https://BioRender.com/mvwxztl.

## Materials and methods

### Cell lines and reagents

All human CRC cell lines (HCT116, SW480, HT29 and RKO) were a kind gift from Dr. Dennis Hsu (University of Pittsburgh, PA). The CRC cell lines were cultured in Dulbecco’s Modified Eagle Medium (DMEM) supplemented with 4.5 g/L glucose, 10% fetal bovine serum (FBS), and 1% penicillin-streptomycin in a humidified atmosphere of 5% CO_2_ at 37°C. All CRC cell lines were authenticated by autosomal short tandem repeat (STR) DNA profiling performed at the University of Arizona Genetics Core (UAGC), prior to starting (Supplementary File 2). U2OS-FAP-TRF1, U2OS-FAP-TRF1 DDB2 KO, and U2OS-FAP-TRF1 mNeon-DDB2 cell lines were cultured in DMEM supplemented with 4.5 g/L glucose, 10% FBS, 1% penicillin-streptomycin, and 500 µg/mL G418, as reported previously^3^. RPE-FAP-TRF1 cells were cultured in Dulbecco’s Modified Eagle Medium/Nutrient Mixture F-12 (DMEM/F12, 1:1) containing 2.438 g/L sodium bicarbonate (Gibco), 10% FBS, 1% penicillin-streptomycin, and 500µg/mL G418 (Gibco), as reported previously ^3^. All cell lines were routinely tested for mycoplasma contamination.

Oxaliplatin (Cat# 09512) and cisplatin (Cat# 232120) were obtained from Sigma-Aldrich (St. Louis, MO). A 2.5 mM stock solution of oxaliplatin was prepared in water by incubating at 37°C for 4 h with occasional vortexing. Cisplatin stock (2.5 mM) was prepared by dissolving in 0.9% NaCl and incubating at 37°C for 2 h with occasional vortexing. Both oxaliplatin and cisplatin stocks were aliquoted and stored at 4°C. Actinomycin D (Cat# A1410) was purchased from Sigma-Aldrich, dissolved in DMSO, and stored at −80°C. Pevonedistat (Cat# HY-70062) and CSN5i-3 (Cat# HY-112134) were procured from MedChemExpress (NJ, USA), dissolved in DMSO, aliquoted, and stored at −80°C.

### Cell viability assays

For all cell viability experiments, 4,000 cells were seeded in each well of a 96-well plate and incubated in a humidified atmosphere of 5% CO_2_ at 37°C overnight. The next day, cells were treated with oxaliplatin or cisplatin for 2 h; after 2 h, the medium was replaced with fresh drug-free medium, and cells were incubated for a further 72 h. For CSN5i-3 and pevonedistat treatment, cells were treated for 72 h. Cell viability was assessed using the CellTiter-Glo 2.0 reagent (Cat# G9242, Promega) according to the manufacturer’s protocol and as previously described ^49^. IC_50_ values were calculated using GraphPad Prism software (version 11).

### Clonogenic assay

For the clonogenic assay, cells were plated in 6-well plates 24 h to treatment. The next day, cells were treated with oxaliplatin or cisplatin for 2 h, after which the medium was replaced with fresh drug-free medium. For CSN5i-3 and pevonedistat, cells were treated for 24 h. Cells were then allowed to grow for 8-10 days. Colonies were fixed with 4% paraformaldehyde (PFA) in PBS for 15 min at room temperature and stained with 0.1% crystal violet in 20% methanol for 30 min at room temperature. Plates were washed with water and dried overnight, and colonies were counted manually.

### Drug combination and synergy analysis

Drug interactions between oxaliplatin and either NAEi (pevonedistat) or CSN5i-3 in HCT116 cells were quantified using the Chou-Talalay median-effect method ^40, 41^. Cells were treated with each agent alone or in fixed-ratio combination at three ascending dose levels (C1, C2, C3), and viability was measured by the CellTiter-Glo assay after 48 and 72 h as described above. Normalized dose-response data for the single agents and the corresponding combinations were entered into CompuSyn software (ComboSyn, Inc.; freely available at https://www.combosyn.com/), which calculates the combination index (CI) as a function of the fraction of cells affected (Fa) at each dose based on the median-effect principle. CI values of <1, =1, and >1 indicate synergistic, additive, and antagonistic interactions, respectively. Fa-CI plots were generated across the full range of Fa (0 - 1) to assess the magnitude and consistency of synergy for each drug combination.

### Gene expression analysis

#### TCGA data-based analysis

Gene expression data and corresponding clinical information were obtained from The Cancer Genome Atlas (TCGA) database (https://portal.gdc.cancer.gov/). RNA-seq gene expression data, along with associated clinicopathological information, were downloaded for the colorectal cancer cohort (TCGA-COAD). Kaplan-Meier survival curves were estimated to evaluate overall survival across the entire colorectal cancer cohort and within the subgroup of stage IV patients treated with oxaliplatin.

#### FOLFOX/oxaliplatin responder vs. non-responder analysis

Transcriptomic datasets and corresponding clinical metadata for a cohort of patients with advanced or metastatic CRC who received first-line combination therapy with FOLFOX/oxaliplatin were obtained from the NCBI Gene Expression Omnibus (GEO) database under accession numbers GSE83129, GSE28702, and GSE69657. The finalized study cohort comprised a total sample size of N = 86 patients. Clinical therapeutic response profiles were evaluated by investigators using the Response Evaluation Criteria in Solid Tumors (RECIST v1.1). Patients documented as achieving a Complete Response (CR) or Partial Response (PR) were classified as Responders (n = 45). Patients exhibiting Stable Disease (SD) or Progressive Disease (PD) were classified as Non-Responders (n = 41). Confounding baseline clinical covariates, including age at diagnosis, tumor pathological stage (III vs. IV), and microsatellite status (microsatellite stable [MSS] vs. microsatellite instability-high [MSI-H]) were compiled directly from the clinical metadata matrices for subsequent multivariate modeling.

#### RNA-seq alignment and transcript quantification

Raw sequencing reads were processed through standard quality control and expression quantification architectures ^42–44, 50^. High-quality paired-end sequence files were mapped to the human reference genome GRCh38 using a splice-aware aligner ^42–44, 50^. Gene-level raw counts were derived using HTSeq-count. To minimize variations in depth and library composition across sequencing runs, the raw genomic matrices were normalized using DESeq2’s median-of-ratios method to account for differences in sequencing depth and library composition. Targeted downstream gene analysis focused specifically on the transcript abundance of the *DDB2* locus.

#### Statistical framework and survival modeling

Baseline variations in continuous, normalized *DDB2* gene expression levels between the stratified chemotherapeutic response groups were evaluated using a two-tailed, unpaired Student’s *t*-test with Welch’s correction to accommodate potential heteroscedasticity between the cohorts. The continuous distribution properties of the expression matrix were confirmed to be normally distributed using a Shapiro-Wilk test (P>0.05). To determine whether *DDB2* transcript expression operated as an independent prognostic factor for patient progression-free survival (PFS), we performed a multivariate Cox proportional hazards regression analysis. The survival model adjusted for continuous *DDB2* expression as an independent variable alongside fixed clinical covariates, specifically age, pathological tumor stage, and microsatellite status. Proportional hazards assumptions for the global model and individual covariates were verified using Schoenfeld residuals. Statistical operations were executed in the R v4.3.1 (with survival package) / python v3.10 (with Lifelines) environment. Structural design of the multivariate Cox proportional hazards model, data organization matrices, and publication-ready code generation were assisted by Google AI (Large Language Model, 2026), with all final statistical metrics, calculations, and mathematical models manually verified and validated by the authors.

### Western blot

For western blot analysis, protein was isolated from the respective cells using 1X lysis buffer as previously reported ^3^. Protein was quantified, and western blots were prepared and performed as previously described ^3^.The following primary antibodies were used: DDB2 (1:1000; Abcam #ab181136; diluted in 10% skimmed milk in 1X PBST [phosphate-buffered saline containing 0.1% Tween-20]), XPC (1:1000; CST #12701S; diluted in 5% skimmed milk in 1X PBST), p21 (1:5000; Abcam #ab109199; diluted in 10% skimmed milk in 1X PBST), β-actin (1:30,000; Sigma #A2228; diluted in 10% skimmed milk in 1X PBST), p27 (1:5000; Abcam #ab32034; diluted in 10% skimmed milk in 1X PBST), and p53 (1:500; Santa Cruz #sc-126; diluted in 5% skimmed milk in 1X PBST). All primary antibodies were incubated overnight at 4°C. The following secondary antibodies were used: anti-rabbit IgG (1:50,000; Sigma #A0545) or anti-mouse IgG (1:50,000; Sigma #A4416). Secondary antibodies were incubated for 1 h at room temperature. Target proteins were detected by chemiluminescence using SuperSignal West Femto and West Pico PLUS reagents (Thermo Fisher Scientific). Blots were analyzed and quantified using ImageJ software.

### Plasmids

mNG-DDB2 K244E-PAM plasmid was made by Gene Universal Inc. pHcRed-NPM1wt-C1 (plasmid #131818) was purchased from Addgene. HaloTag-JF650-DDB2, HaloTag-JF554-DDB2, and DDB1-eGFP were acquired from a previous study ^28^.

### Transfections and nuclear extraction

For confocal experiments, 0.25 µg of either pHcRed-NPM1 plasmid or mNG-DDB2 K244E-PAM plasmid DNA was mixed with 1 µL of Lipofectamine 2000 (Thermo Fisher Scientific, #11668027) in Opti-MEM medium according to the manufacturer’s instructions. Fresh complete media were added 5 h post-transfection. Images were acquired using a confocal microscope 24 h post-transfection. For nuclear extracts, U2OS cells were transfected with 4 µg of plasmid DNA containing fluorescent-tagged proteins of interest, as reported previously. ^28^. Cells overexpressing HaloTag fusion proteins were labeled with 100 nM Janelia Fluor® JF650 (HT1070, Promega) or JF554 (HT1030, Promega) HaloTag ligand for 30 min at 37 °C. Nuclear extraction was performed using a nuclear extraction kit (Abcam, ab113474), as previously described.^28^.

### Lambda DNA substrate generation

Lambda DNA was biotinylated and isolated for C-trap experiments as previously described ^28^, aliquoted at 20 ng/µL, and stored at −20 °C. After thawing, aliquots were stored at 4 °C for up to 2 weeks and then discarded. One microgram of lambda DNA (50 µL at 20 ng/µL) was dialyzed against 500 mL of TE buffer (10 mM Tris, pH 8.0, 1 mM EDTA) at 4 °C with continuous stirring; the buffer was exchanged twice, once after 24 h and again after 12 h. The volume and concentration of the dialyzed DNA were then measured. Ninety nanograms of dialyzed lambda DNA (18 µL at 5 ng/µL) were incubated with 2 µL of either 37.5 µM cisplatin or 250 µM oxaliplatin (final concentrations of 3.75 µM and 25 µM, respectively) in a PCR tube in the dark at 37 °C for 2 h. Reactions were brought to room temperature and quenched with 1 µL of 4 M NaCl (final concentration, 0.2 M), then dialyzed overnight against 500 mL of TE buffer containing 0.1 M NaCl at 4 °C with continuous stirring. The volume and concentration of the dialyzed DNA were measured again. Twenty nanograms of each cisplatin- or oxaliplatin-treated lambda DNA were mixed with 500 µL of PBS and used for C-trap experiments.

### Design of cisplatin- and oxaliplatin damaged oligonucleotides for single-molecule analysis

#### Synthesis of cisplatin and oxaliplatin intrastrand crosslink oligonucleotides

Cisplatin and oxaliplatin oligonucleotides were prepared as described. ^51^. In a labeled Eppendorf tube, 20 nmol of oligonucleotide was diluted to 600 μL at a final concentration of 10mM sodium perchlorate (NaClO_4_) and 5mM acetic acid and treated with 1 equivalent of activated cisplatin or 3 equivalents of oxaliplatin for 1 hour at 37 °C, protected from light. The sequence of the oligonucleotides was as follows: for 1,2-intrastrand adducts: 5’-Phos-TCT TCT TCT **GG** TAC TCT TCT TCT-3’; oligonucleotides for 1,3-intrastrand adducts: 5’-Phos-TCT TCT TCT **G**T**G** CAC TCT TCT TCT-3’. After incubation, excess cisplatin or oxaliplatin was removed using a 0.2 μm nylon filter. The oligonuelcotides were purified using an AKTA Pure FPLC as follows; A MonoQ 5/50 GL column was operated using a gradient of (A) 10mM NaOH and (B) 1M NaCl in 10mM NaOH at 2 mL/min starting at 10% B for 10 CV to equilibrate, then kept at 10% B for 5 CV after sample injection, followed by increasing to 30% B over 5 CV, then to 50% B over 40 CV, then to 100% B over 20 CV, held constant for an additional 5 CV and then decreased to 10% B to re-equilibrate for 10 CV. Under these conditions, the 23mer 1,2- and 24mer 1,3-intrastrand crosslink substrate eluted between 62 – 65 CV, and the un-platinated oligo was not observed. All collected substrates were dried using lyophilization. The resulting solid was resuspended in 1 mL of UP-water and desalted using centrifugal filtration (Merck, Amicon® Ultra 10 kDa filters) at 14,000 rcf for 10 minutes at 4 °C. The sample was washed with UP-water an additional three times to ensure buffer exchange. The product was confirmed by negative mode full-scan LC-MS.

#### Site-specific oligonucleotide substrate generation for C-trap

Synthetic DNA oligonucleotides (Integrated DNA Technologies) were designed to generate a fluorescein-labeled thymine nicked duplex substrate. The upper left strand contained a fluorescein fluorophore (/5phos/CAACCAGCTGAACATGCCTTTTGA/iFluorT/TGAGCAG-3′), the upper right strand was 5′ phosphorylated, 5′-Phos-TCTTCTTCT**GG**TACTCTTCTTCT-3′; (for 1,2 GG) and 5′-Phos-TCTTCTTCT**G**T**G**CACTCTTCTTCT-3′; (for 1,3-GTG), and the complementary lower strands were 3′GTCGACTTGTACGGAAAACTAACTCGTC AGAAGAAGACCATGAGAAGAAGAACCA 5’ phos and 3′ GTCGACTTGTACGGAAAACTAACTCGTC AGAAGAAGACACGTGAGAAGAAGAACCA 5’ phos respectively.

Equimolar concentrations (1 µL) of all three oligonucleotides were mixed with 17 µL of annealing buffer (100 mM KCl, 10mM Tris-HCl pH 8.0). The reaction mixture was heated to 95 °C for 5 min, cooled to RT. It was then treated with 400 units of T4 ligase (∼200 fold excess) in 1X PNK buffer and incubated overnight at RT. It was then heat-inactivated at 65 °C for 15 min, followed by slow cooling, and then 2 µL aliquots were prepared and stored at −20 °C for further use.

### Arms ligation for C-trap

Using the LUMICKS DNA tethering kit and protocol, a sequence of interest was ligated into two handles that are each 6.3 kb in length to generate a substrate that is 12.6 kb in total. Briefly, in a 10 μL reaction, 25 nM of the duplex, 2.5 μL of each handle, 1 μL of 10X ligase buffer, and 0.5 μL of ligase were mixed and incubated at room temperature overnight in dark. The reaction was heat-inactivated at 65 °C for 10min and allowed to slowly cool to room temperature. Substrate was stored at 4 °C and protected from light.

### Single-molecule experiments

Single-molecule experiments were performed on a LUMICKS C-trap using a 5-channel flow chamber as previously described ^28, 30, 52, 53^. Briefly, for experiments for the platinum treated λ DNA (∼48.5 kb), channels one, two, and three were filled with 4.0-4.9 μm polystyrene streptavidin beads (LUMICKS), biotinylated platinum treated DNA, and UV-DDB buffer containing 20mM HEPES pH7.5,150mM NaCl, 0.5 mg/ml BSA,1mM DTT, 1 mMTrolox, and 5% glycerol and the Halo-tagged JF-650 /JF-554 labelled UV-DDB2 nuclear extracts were diluted 1:10 ratio in the same buffer and flowed in channel 4/5. For the 12 kb substrates, 1.5-1.7 μm streptavidin beads were utilized using the same buffer and extract conditions as described above. The data were exported with Bluelake and analyzed using custom software from LUMICKS (Pylake) as previously reported ^28, 30, 52, 53^.

### Fixed cell confocal imaging

For U2OS-FAP-TRF1 cells stably expressing mNeonGreen-DDB2, 200,000 cells were seeded onto coverslips (Fisher Scientific, Cat# 12-544-10; 22 X 22 mm, No. 1) in a 6-well plate overnight prior to transfection with the NPM1 vector (pHcRed-NPM1wt-C1) (Addgene plasmid #131818). For U2OS-FAP-TRF1-DDB2 KO cells, cells were co-transfected with the NPM1 and mNG-DDB2 K244E-PAM plasmids (Gene Universal). After 24 h, cells were treated with the respective drugs for 6 h. Following treatment, cells were washed three times with phosphate-buffered saline (PBS) and fixed with 4% PFA for 20 min at room temperature. Cells were then washed three times with PBS, stained with DAPI (0.5 µg/mL in PBS) for 10 min at room temperature, washed three more times with PBS, and mounted onto slides using ProLong Diamond Antifade Mountant (#P36970; Molecular Probes).

Images were acquired on a Nikon Eclipse Ti inverted confocal microscope using a Plan Apo VC 60X oil-immersion objective (DIC N2). Fluorescence excitation wavelengths were 488 nm for mNeonGreen-DDB2, 405 nm for DAPI, and 561 nm for pHcRed-NPM1. Fluorescence emission was collected at 525 nm for mNeonGreen-DDB2, 450 nm for DAPI, and 595 nm for pHcRed-NPM1, using the standard instrument detector. The pinhole diameter was set to 24.27 µm. All settings were kept the same across samples and throughout the experiment. Images were saved in ND2 file format and analyzed using NIS-Elements software (Nikon, version 5.2.1).

### Live-cell confocal imaging

Confocal fluorescence images were acquired on a Nikon Eclipse Ti inverted microscope equipped with a Nikon A1 confocal laser-scanning head and a 60X 1.40 NA objective. All experiments were carried out at 37°C in a pre-mixed atmosphere of 5% CO_2_, 20% O_2_, balanced with nitrogen, using a Tokai Hit live-cell incubation chamber. A total of 100,000 U2OS-FAP-TRF1 cells stably expressing mNeonGreen-DDB2 were seeded in a 35 mm glass-bottom dish (Cat# P35G-1.5-14-C; MatTek Life Sciences) 24 h prior to transfection. After 24 h, cells were transiently transfected with the pHcRed-NPM1 plasmid. At 24 h post-transfection, drugs were added immediately before imaging. Images were captured for 6 h at 4 images per hour (1 image every 15 min). Fluorescence excitation and emission settings were the same as those used for fixed-cell imaging, using the standard instrument detector. Sequential images were acquired as 12-bit, two-channel images at a fixed pixel size of 0.21 µm, with the pinhole diameter set to 1.0 Airy unit, within a 1024 X 1024-pixel region of interest centered on the cell. A pixel dwell time of 1.1 µs was used, resulting in a frame time of 100 ms. Images were processed using NIS-Elements software (Nikon, version 5.2.1).

### Immunohistochemistry (IHC)

For IHC, unstained formalin-fixed paraffin-embedded (FFPE) colon cancer tissue microarrays (TMAs) were procured from TissueArray.com (Cat# CRM1503 and Cat# CRC1601CSur). The TMA slides were baked for at least 1 h in an oven at 60°C prior to immunostaining. Staining was performed as per the manufacturer’s instructions (Proteintech, Cat# PK10017). Briefly, paraffin was removed using xylene, followed by heat-induced antigen retrieval in sodium citrate buffer. Eight to ten drops of blocking buffer (Part No. HC003, Proteintech, Cat# PK10017) were added to each slide for 30 min. Slides were then incubated with DDB2 primary antibody (1:1500; Abcam #ab181136) for 1 h at room temperature. Stained tissues were imaged at 20X magnification. Visual, semi-quantitative scoring was performed by a trained pathologist at TissueArray.com using a simple scoring rubric of 1+ to 3+ staining intensity, with 0+ representing no visible color reaction. In the percentage quantification of DDB2-positive cells in normal versus paired tumor tissue, the cells scored within a range (<25% or >75%) were assigned the midpoint of that range for statistical analysis.

### Statistics and reproducibility

All experimental assays were performed at least in triplicate, with exact replicate numbers indicated in the respective figure legends. Continuous data are presented as mean ± standard error of the mean (s.e.m.). All statistical analyses were performed using GraphPad Prism software (version 11). Statistical significance was determined by unpaired or paired two-tailed Student’s *t*-test, or one-way ANOVA with Bonferroni’s or Tukey’s multiple comparisons test, as appropriate.

## Supporting information

Combine Supplementary Figures

Description of Additional Supplementary Files

Supplementary File 2_STR

Western blot full scan

## Data availability

Further information and requests for reagents and raw kymograph files should be directed to the corresponding author. All kymograph measurements and uncropped blots are available in the Source Data file. Source data are provided with this paper.

## Acknowledgement

The authors thank Matthew A. Schaich for technical assistance in the C-trap experiments and Shahjahan for assistance with the cell viability experiments. We used the online AI tool, Anthropic Claude to make the animation for supplementary video 4. The content was reviewed and validated by the authors, who take full responsibility for the content.

## Author contributions

Conception and design: VK and BVH. Data acquisition: VK, AN, AM, VR, KEH, CW, M C, SCW, DH, ORD and BVH. Data analysis and interpretation of data: VK, AN and BVH. Writing (original draft): VK and BVH. Writing (review and editing): VK, AN, OSD, DH and BVH.

## Ethics declaration

Not applicable.

