## Supplementary material for "DDB2 localizes to the nucleolus to mediate oxaliplatin resistance in colorectal cancer": Combine Supplementary Figures

#### Supplementary Fig S1.

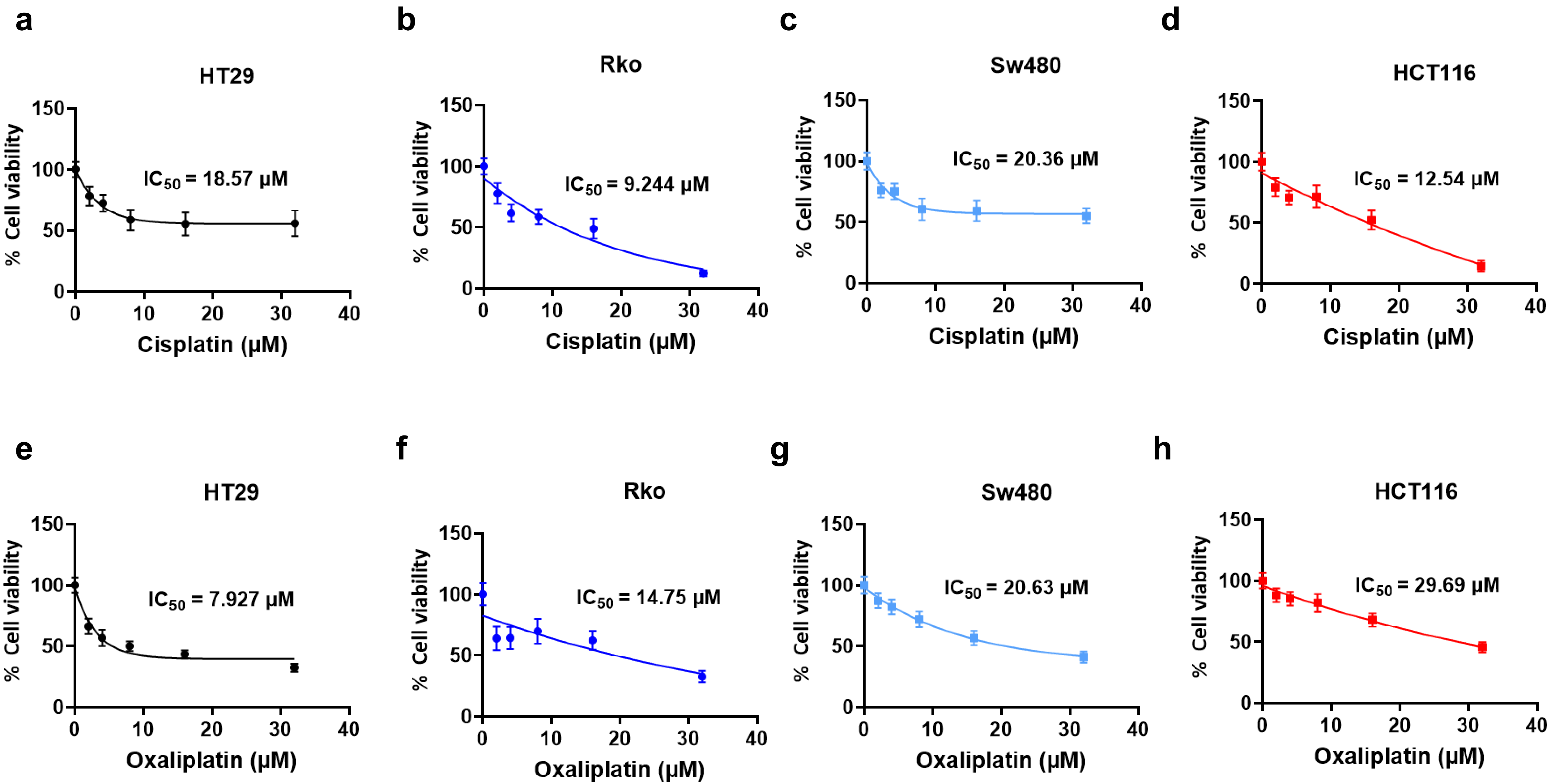

i

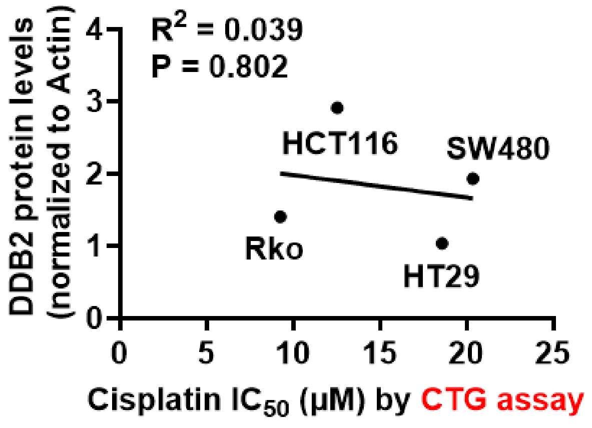


j

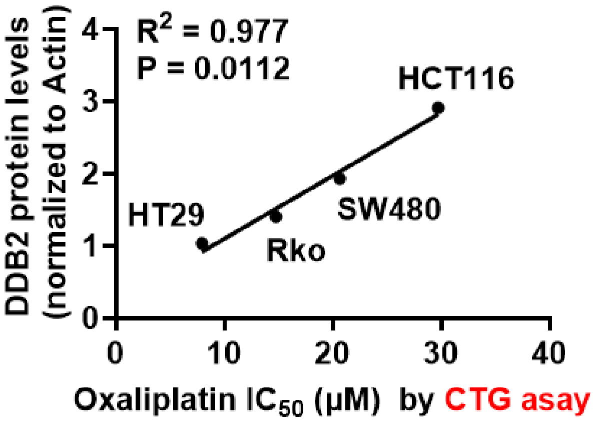


k

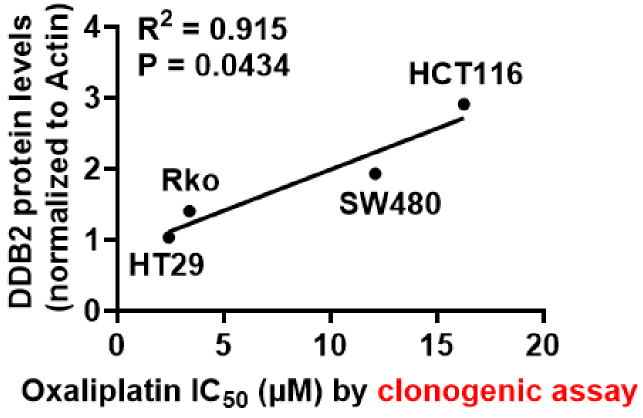


l

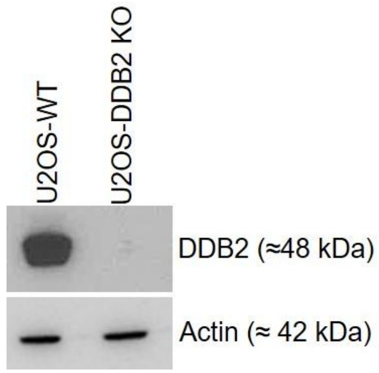


m

| | DDB2-WT | DDB2-KO |
|---|---|---|
| **Cisplatin (μM)** | 5.64 | 3.37 |
| **Oxaliplatin (μM)** | 11.03 | 4.14 |

n

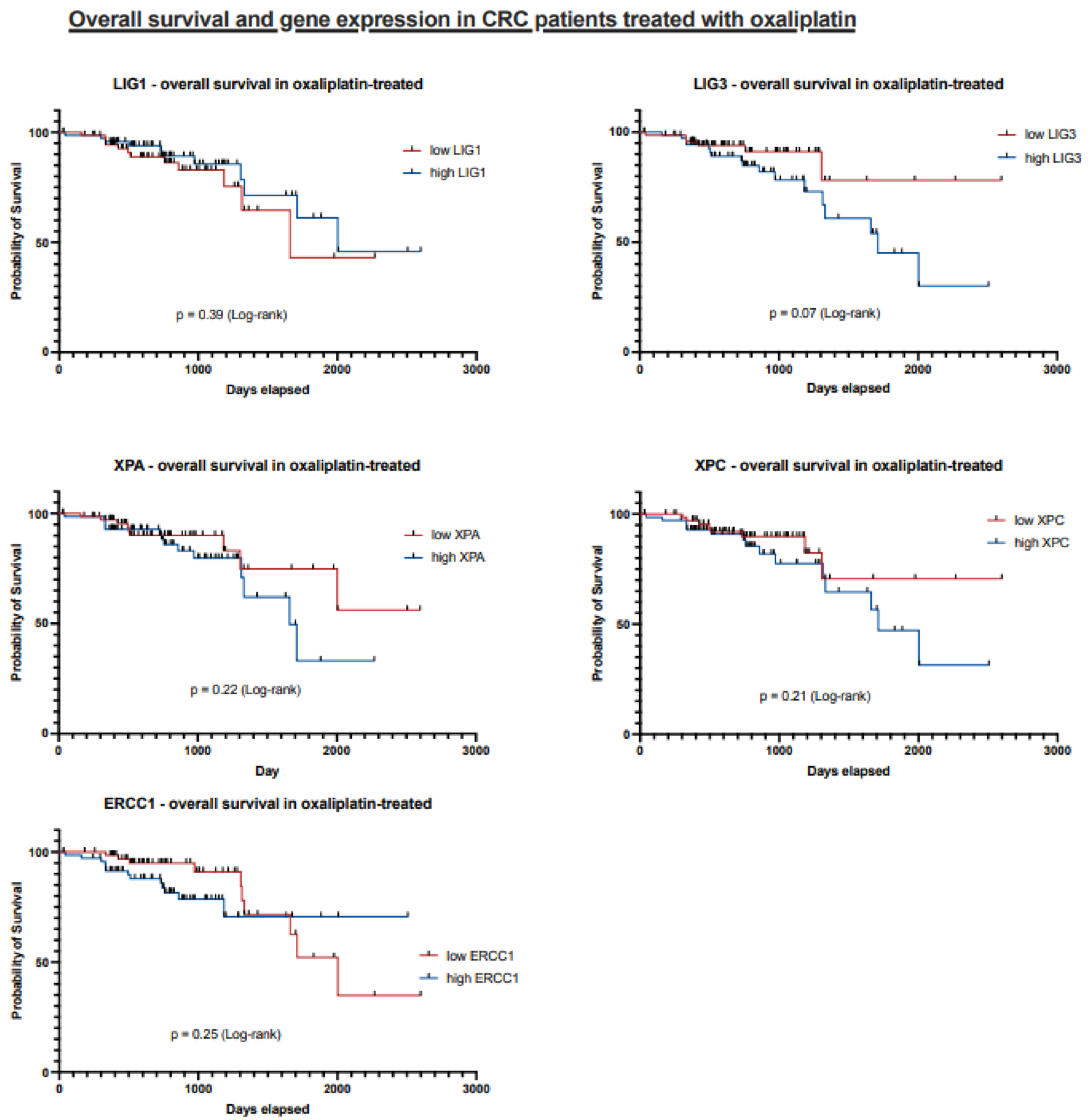
Overall survival and gene expression in CRC patients treated with oxaliplatin
LIG1 - overall survival in oxaliplatin-treated
low LIG1
high LIG1
p = 0.39 (Log-rank)
LIG3 - overall survival in oxaliplatin-treated
low LIG3
high LIG3
p = 0.07 (Log-rank)
XPA - overall survival in oxaliplatin-treated
low XPA
high XPA
p = 0.22 (Log-rank)
XPC - overall survival in oxaliplatin-treated
low XPC
high XPC
p = 0.21 (Log-rank)
ERCC1 - overall survival in oxaliplatin-treated
low ERCC1
high ERCC1
p = 0.25 (Log-rank)
Probability of Survival
Days elapsed
Day
0
50
100
1000
2000
3000

o

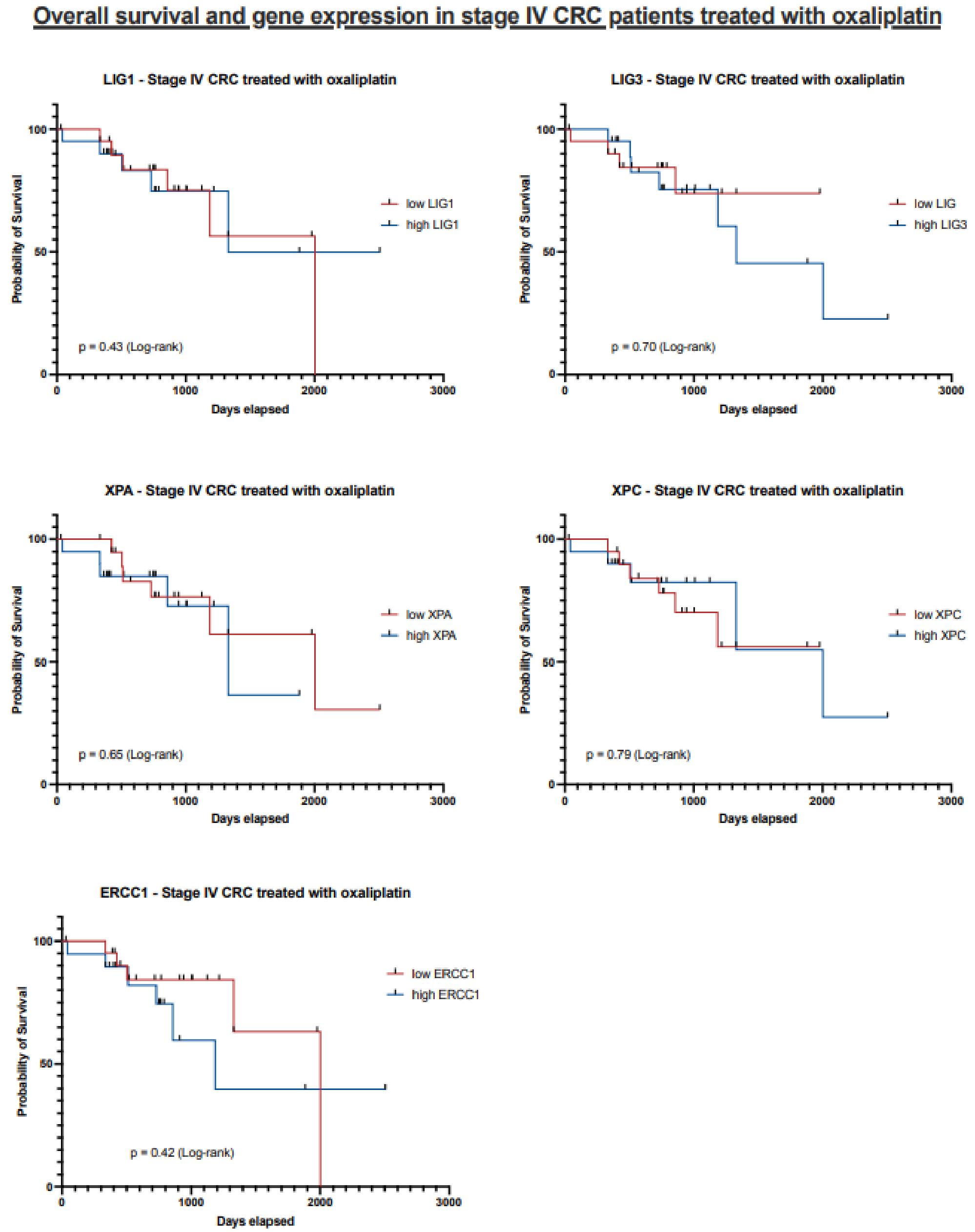
Overall survival and gene expression in stage IV CRC patients treated with oxaliplatin
LIG1 - Stage IV CRC treated with oxaliplatin
low LIG1
high LIG1
p = 0.43 (Log-rank)
LIG3 - Stage IV CRC treated with oxaliplatin
low LIG
high LIG3
p = 0.70 (Log-rank)
XPA - Stage IV CRC treated with oxaliplatin
low XPA
high XPA
p = 0.65 (Log-rank)
XPC - Stage IV CRC treated with oxaliplatin
low XPC
high XPC
p = 0.79 (Log-rank)
ERCC1 - Stage IV CRC treated with oxaliplatin
low ERCC1
high ERCC1
p = 0.42 (Log-rank)
Probability of Survival
Days elapsed
0
50
100
1000
2000
3000

**Supplementary Fig. S1.** Sensitivity of colorectal cancer cell lines to cisplatin and oxaliplatin. **a - d** Dose - response curves for HT29 (**a**), RKO (**b**), SW480 (**c**), and HCT116 (**d**) cells treated with cisplatin. **e - h** Dose - response curves for HT29 (**e**), RKO (**f**), SW480 (**g**), and HCT116 (**h**) cells treated with oxaliplatin. Cells were treated either with cisplatin or oxaliplatin for 2 h, and viability (CellTiter-Glo) was assessed after 72 h. Viability is expressed as a percentage of vehicle - treated control cells. Data are presented as mean ± s. e. m.; n = 10, from two independent experiments. $IC_{50}$ values were calculated by fitting the data to a nonlinear regression model in GraphPad Prism and are shown in each panel. The correlation between DDB2 protein levels and the $IC_{50}$ of cisplatin and oxaliplatin was determined by Pearson regression both for the CTG assay (**i - j**) and clonogenic assay (**k**). **l** A western blot showing the complete KO of DDB2 in U2OS cells. **m** $IC_{50}$ of cisplatin and oxaliplatin in DDB2 WT and DDB2 KO U2OS cells obtained from a clonogenic assay. **n** Kaplan-Meier overall survival curves for all colorectal cancer patients treated with oxaliplatin (TCGA-COAD), stratified by low versus high tumor expression of LIG1, LIG3, XPA, XPC, and ERCC1. **o** Kaplan-Meier overall survival curves for stage IV colorectal cancer patients treated with oxaliplatin only, stratified by low versus high tumor expression of LIG1, LIG3, XPA, XPC, and ERCC1.

#### Supplementary Fig S2.

a

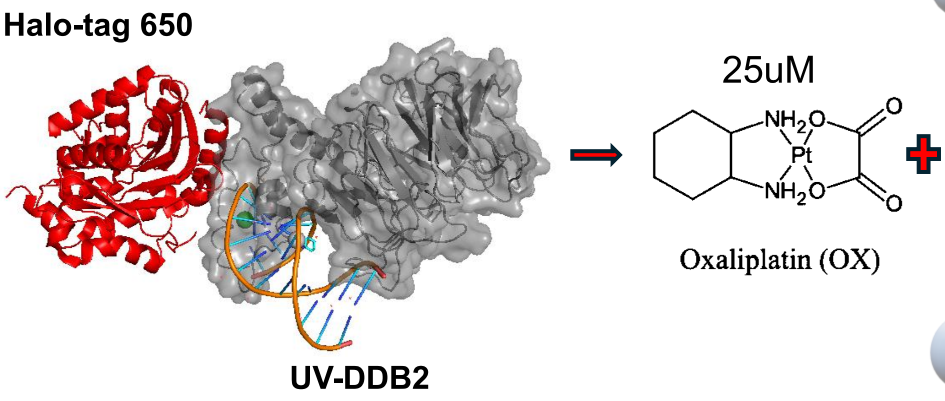


b

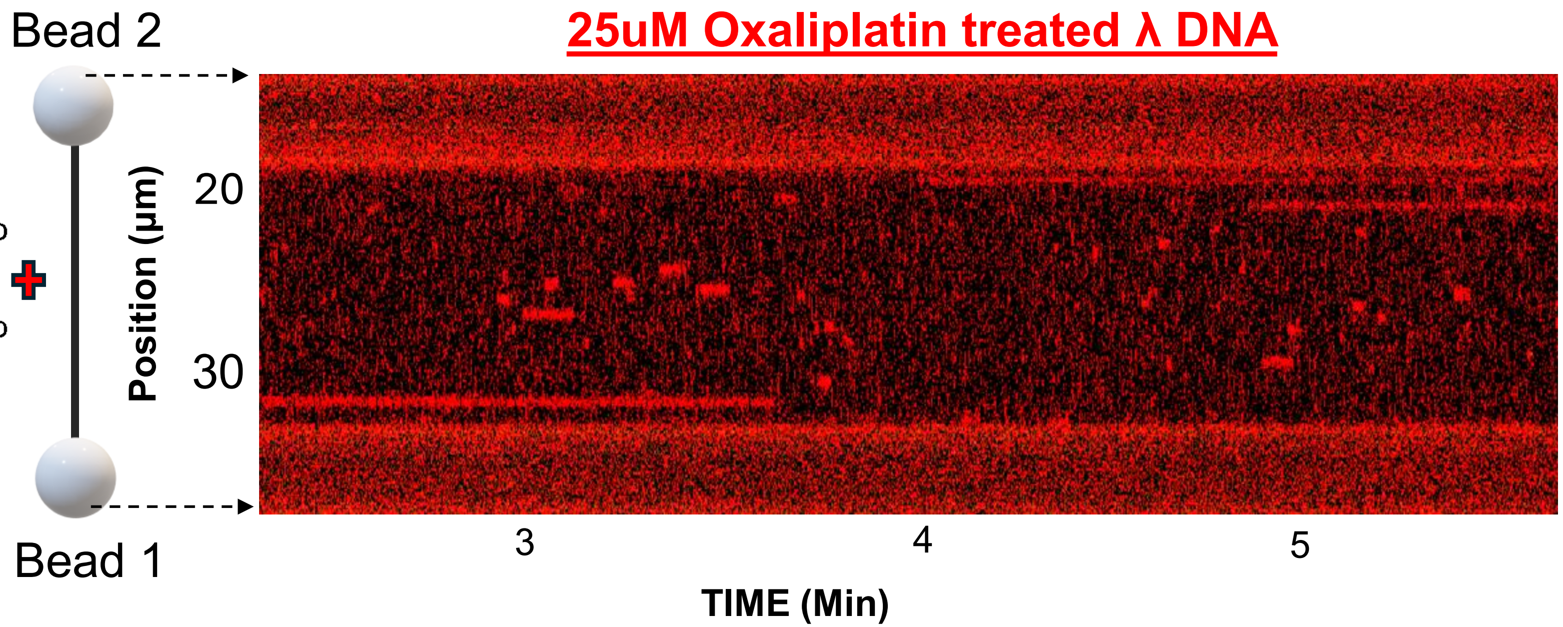


c

Halo-650 DDB2 with 25 uM Oxaliplatin

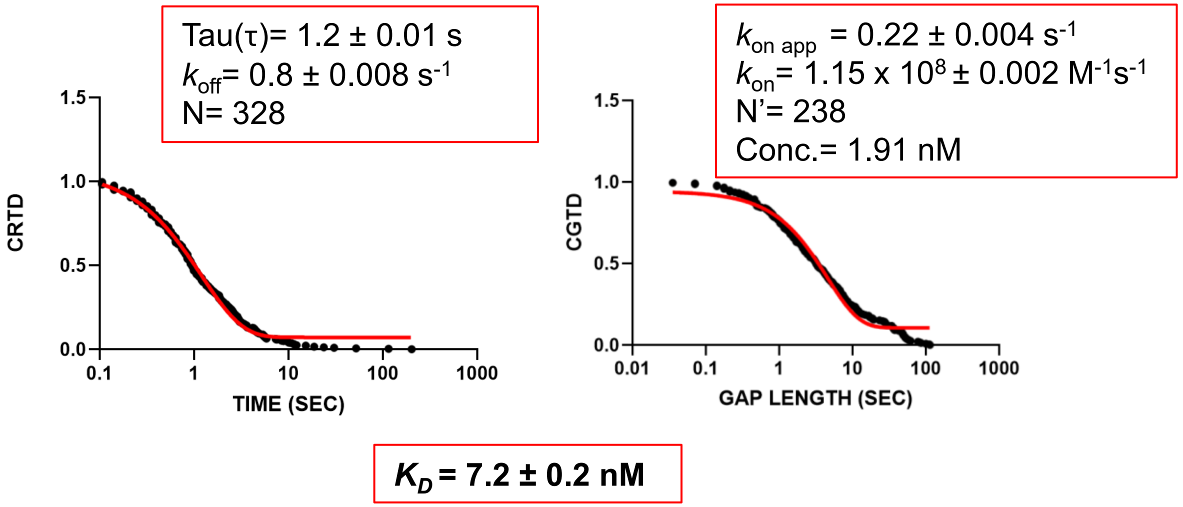

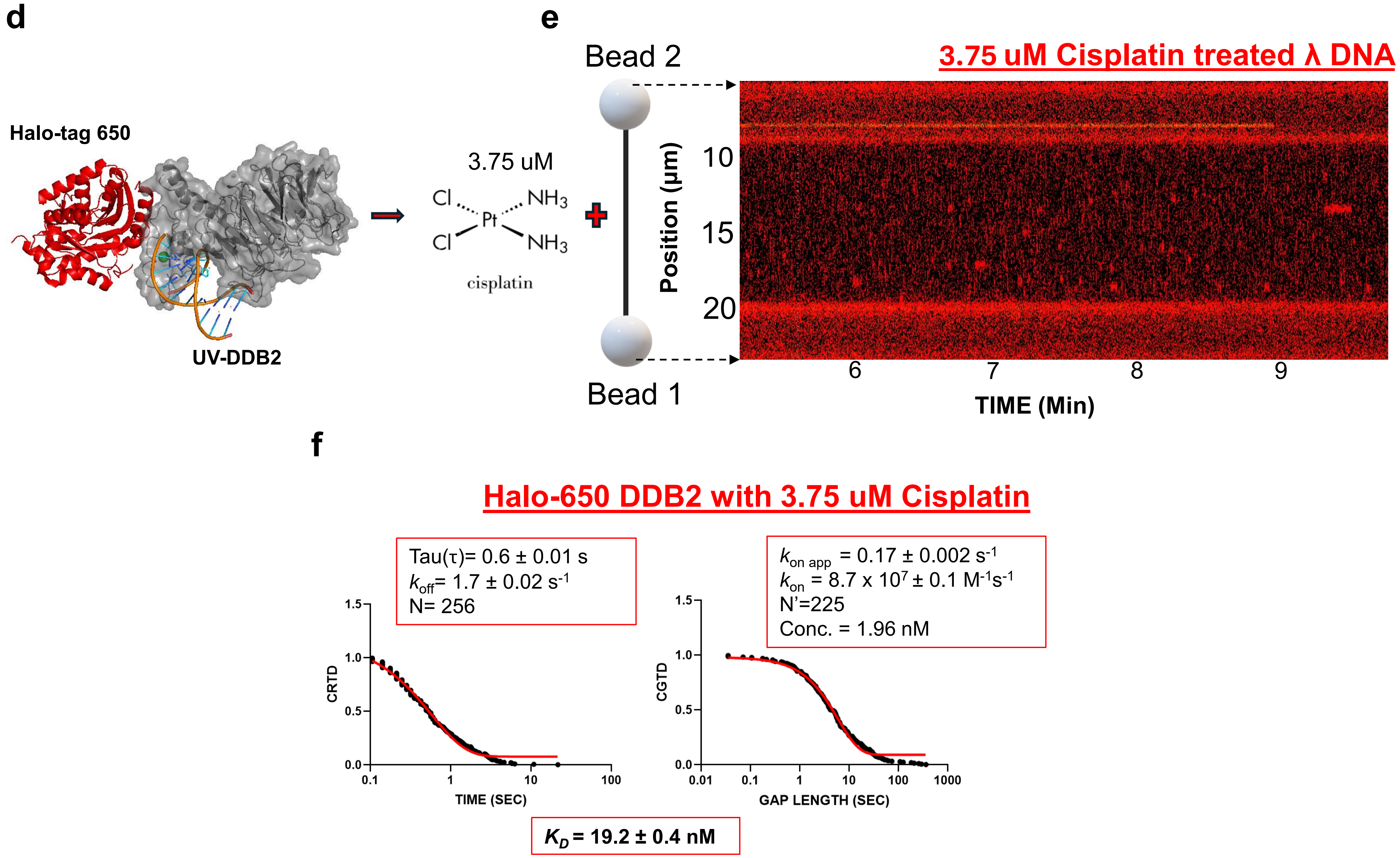
d
Halo-tag 650
UV-DDB2
3.75 uM
Cl
Cl
Pt
NH3
NH3
cisplatin
e
Bead 2
Bead 1
3.75 uM Cisplatin treated λ DNA
Position (μm)
10
15
20
6
7
8
9
TIME (Min)
f
Halo-650 DDB2 with 3.75 uM Cisplatin
Tau(τ)= 0.6 ± 0.01 s
k_off = 1.7 ± 0.02 s^-1
N= 256
k_on app = 0.17 ± 0.002 s^-1
k_on = 8.7 x 10^7 ± 0.1 M^-1 s^-1
N'=225
Conc. = 1.96 nM
CRTD
TIME (SEC)
CGTD
GAP LENGTH (SEC)
1.5
1.0
0.5
0.0
0.1
1
10
100
0.01
1000
K_D = 19.2 ± 0.4 nM

g

COMBINED TABLE OF BINDING AFFINITIES OF HALO-UV-DDB2 TO PLATIN TREATED λ DNA

| PLATIN LESION | Lifetime (s) | $k_{off}$ ($s^{-1}$) | N | $K_{on\ app}$ ($s^{-1}$) | $k_{on}$ ($M^{-1}s^{-1}$) | N' | $K_D$ (nM) | Conc. (nM) |
|---|---|---|---|---|---|---|---|---|
| **25uM Oxaliplatin treated λ DNA** | 1.2 ± 0.03 | 0.8 ± 0.008 | 328 | 0.22 ± 0.004 | $1.15 \times 10^{8} \pm 0.002$ | 238 | 7.2 ± 0.2 | 1.91 |
| **3.75 uM Cisplatin treated λ DNA** | 0.6 ± 0.01 | 1.7 ± 0.02 | 256 | 0.17 ± 0.002 | $8.7 \times 10^{7} \pm 0.1$ | 225 | 19.2 ± 0.4 nM | 1.96 |

**Supplementary Fig S2. Single-molecule analysis of Halo-tag 650 - DDB2 binding to oxaliplatin- and cisplatin-damaged λ DNA lesions. a** Representative structural model of N-terminal Halo-tag 650 - DDB2 bound to oxaliplatin damaged DNA . Halo - tag 650 is shown in red and DDB2 in gray. **b** Schematic of the C-trap assay and representative kymograph showing Halo-tag 650-DDB2 binding to an oxaliplatin-treated lambda DNA tethered between two optically trapped beads. **c** Cumulative residence time (CRTD) and cumulative gap time (CGTD) distributions for DDB2 binding to oxaliplatin treated lambda DNA. Single-exponential fits are shown in red, and the corresponding binding lifetime, kinetic parameters, and $K_D$ values are indicated in each panel. **d** Representative structural model of N-terminal Halo-tag 650-DDB2 bound to cisplatin-damaged DNA. Halo - tag 650 is shown in red and DDB2 in gray. **e** Schematic of the C - trap assay and representative kymograph showing Halo-tag 650 - DDB2 binding to a cisplatin-treated lambda DNA tethered between two optically trapped beads. **f** Cumulative residence time (CRTD) and cumulative gap time (CGTD) distributions for DDB2 binding to cisplatin-treated lambda DNA. Single-exponential fits are shown in red, and the corresponding binding lifetime, kinetic parameters, and $K_D$ values are indicated in each panel. **g** Combined table of binding affinities of Halo-DDB2 to platinum-treated λ DNA.

### Supplementary Fig S3.

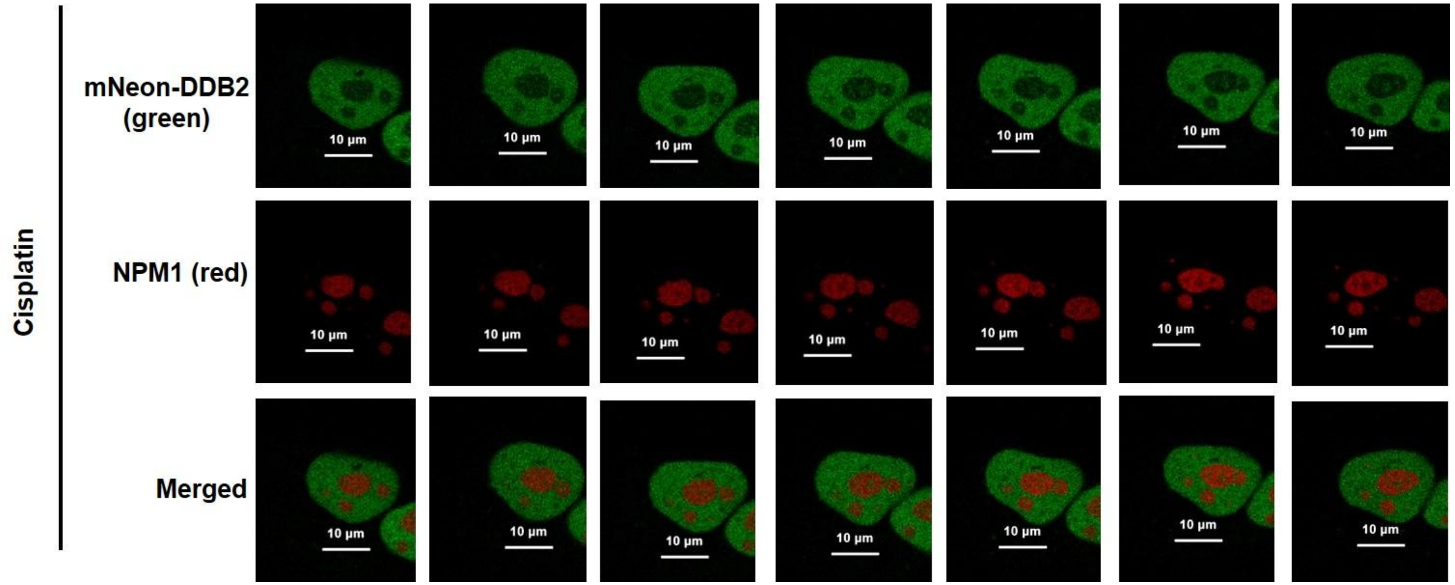


**Supplementary Fig S3.** Time-lapse live-cell imaging of mNeonGreen-DDB2 (green) and NPM1-pHcRed (red) in cisplatin-treated cells, imaged every 15 min for 6 h (representative frames shown at 1 h intervals). Corresponding time-lapse movies are provided in Supplementary Video 2.

#### Supplementary Fig S4.

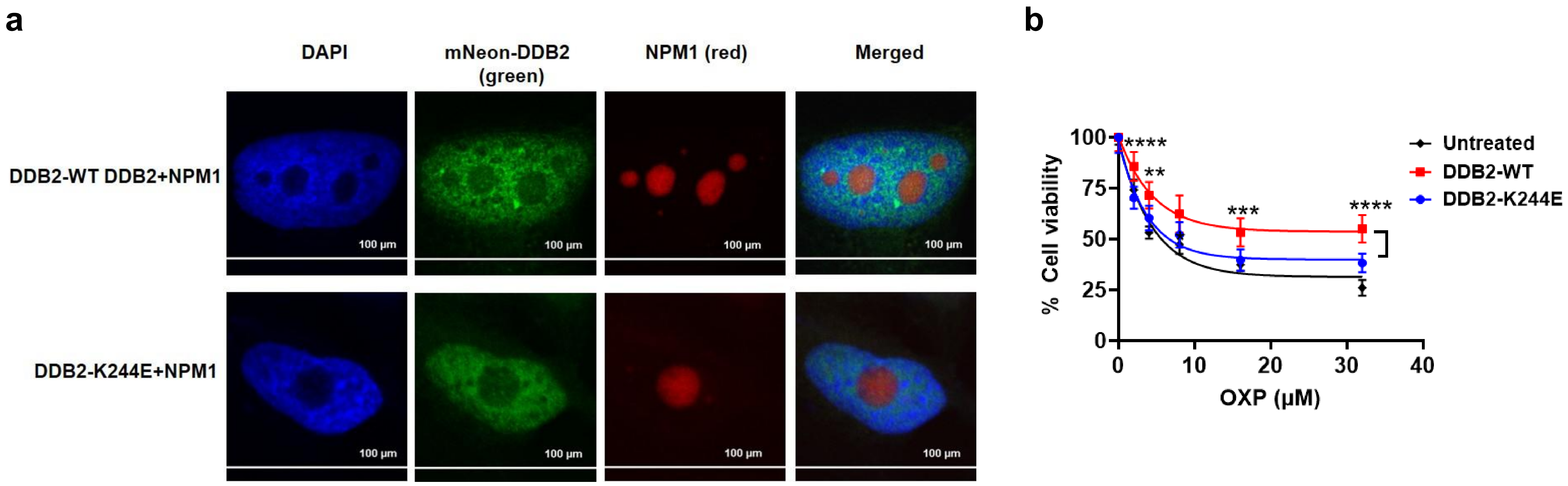


**Supplementary Fig S4 a** Representative confocal images of U2OS DDB2-KO cells co-expressing mNeonGreen-DDB2-WT or DDB2-K244E variant and NPM1-pHcRed, showing DAPI (nuclear DNA, blue), mNeon-DDB2 (green), NPM1 (red), and merged channels in untreated cells. Scale bars, 100 µm. **b** Cell viability (CellTiter-Glo; CTG) of the same cell populations following oxaliplatin treatment, shown as % viability relative to untreated controls. The cells were treated with oxaliplatin for 2h and CTG readings were taken after 72 h. Data are mean ± s.e.m. from n = 8, from two independent experiments. **P < 0.01, ***P < 0.001, ****P < 0.0001 by two-way ANOVA with Sidak's multiple comparisons test, DDB2-WT vs. DDB2-K244E.

#### Supplementary Fig S5.

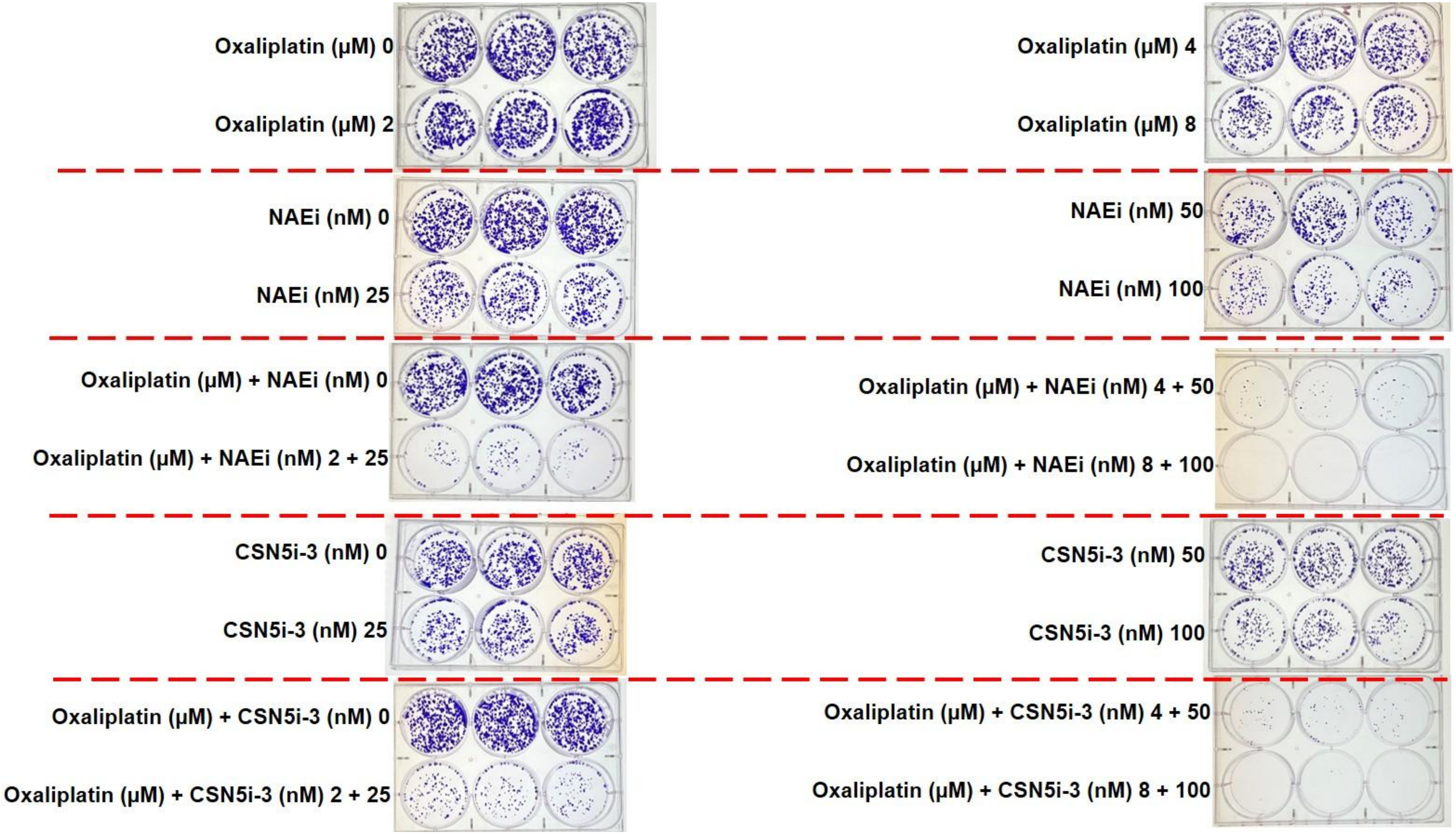

**Supplementary Fig S5. Pharmacological inhibition of DDB2 activity synergizes with oxaliplatin in colorectal cancer cells.** Clonogenic assay images of HCT116 cells treated with oxaliplatin, NAEi, or CSN5i-3 alone, or the indicated combinations, stained with crystal violet. n = 3.

### Supplementary Fig S6.

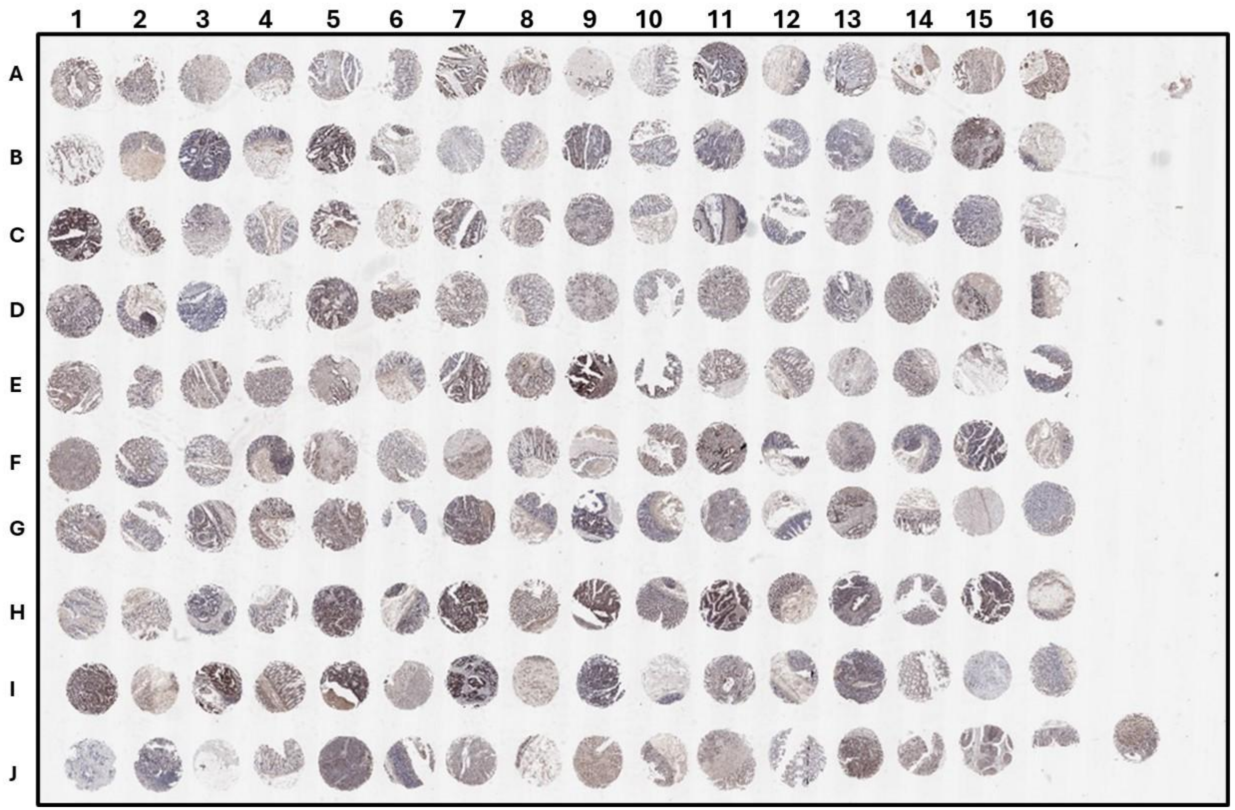


**Supplementary Fig S6. DDB2 protein expression across a colorectal cancer tissue microarray (TMA).**

Immunohistochemical (IHC) staining for DDB2 was performed on a commercially available colorectal cancer tissue microarray (TMA; Cat# CRC1601CSur, Tissue Array.com), containing formalin-fixed, paraffin-embedded tissue cores arranged in a 16 × 10 grid format (columns 1-16, rows A-J). Representative brightfield images show variable DDB2 staining intensity across individual tumor cores, reflecting heterogeneous DDB2 expression among colorectal cancer patient samples.
