## Supplementary material for "DDB2 localizes to the nucleolus to mediate oxaliplatin resistance in colorectal cancer": Description of Additional Supplementary Files

**Title: Supplementary Video 1-3.**

U2OS cells were treated with the IC₅₀ of each drug. Cells were imaged by live-cell confocal microscopy at 37 °C and 5% CO₂, with one image acquired every 15 min for 6 h, beginning at drug addition for 6 hours (25 frames per video). For better visibility, readers are advised to view the videos at 0.25 X playback speed (via the video player settings).

**Supplemental video 1:** Time-lapse live-cell imaging of mNeonGreen-DDB2 and NPM1-pHcRed transfected sham-treated U2OS cells.

**Supplemental video 2:** Time-lapse live-cell imaging of mNeonGreen-DDB2 and NPM1-pHcRed-transfected U2OS cells treated with 5.64 µM cisplatin.

**Supplementary Video 3:** Time-lapse live-cell imaging of mNeonGreen-DDB2 and NPM1-pHcRed-transfected U2OS cells treated with 11.03 µM oxaliplatin.

**Title: Supplementary Video 4.**

**Animated summary of the proposed working model:**  1) DDB2 binds both cisplatin and oxaliplatin DNA lesions. 2) Oxaliplatin, but not cisplatin treatment, drives DDB2 into the nucleolus. 3) The repair-deficient K244E DDB2 variant does not protect cells from oxaliplatin. 4) High DDB2 levels are associated with oxaliplatin resistance in patients, and DDB2 degradation is proposed to restore response. The animation was generated using the online tool Anthropic Claude.
