## Supplementary File 2_STR for "DDB2 localizes to the nucleolus to mediate oxaliplatin resistance in colorectal cancer"

### Cell Line Authentication Report

**Prepared for:** Vinod Kumar
Van Houten, Bennett (PITT) Lab

**Submission:** UAGC-VK-8188809


UNIVERSITY OF ARIZONA GENETICS CORE

University of Arizona Genetics Core
1657 E Helen St Rm 111
Tucson , AZ
85721

http://uagc.arl.arizona.edu


UAGC's comments about sample match is based on the reference data available at the time of comparison.

**Reference STRs**

| Sample Name | D5S818 | D13S317 | D7S820 | D16S539 | VWA | TH01 | AM | TPOX | CSF1PO | Multiple Profiles Seen | Match Comments |
|---|---|---|---|---|---|---|---|---|---|---|---|
| C1R | 11<br>13 | 8<br>11 | 8<br>10 | 11<br>13 | 16<br>17<br>22 | 6<br>10 | X<br>X | 11<br>11 | 8<br>10 | NO | No reference profile found. Sample matches RKO above the 80% match threshold. This is considered a match to RKO (GNE). |
| C2S8 | 13<br>13 | 12<br>12 | 8<br>8 | 13<br>13 | 16<br>16 | 8<br>8 | X<br>X | 11<br>11 | 13<br>14 | NO | No reference profile found. Sample matches SW 480 above the 80% matching threshold. This is considered a match to SW 480 (GNE). |
| C3H6 | 10<br>11 | 10<br>12 | 11<br>12 | 11<br>13 | 17<br>22 | 8<br>9 | X<br>X | 8<br>9 | 7<br>10 | NO | No reference profile found. Sample matches HCT 116 above the 80% matching threshold. This is considered a match to HCT 116 (ATCC). |
| C4C5 | 10<br>13 | 10<br>12 | 9<br>10 | 12<br>13 | 15<br>15 | 8<br>9 | X<br>X | 11<br>11 | 11<br>11 | NO | No reference profile found. Sample matches COLO 205 above the 80% matching threshold. This is considered a match to COLO 205 (GNE). |
| C5H9 | 11<br>12 | 11<br>12 | 10<br>10 | 11<br>12 | 17<br>19 | 6<br>9 | X<br>X | 8<br>9 | 12<br>12 | NO | No reference profile found. Sample matches HT-29 above the 80% matching threshold. This is considered a match to HT-29 (GNE). |

**Additional STRs**

| Sample Name | D3S1358 | D21S11 | D18S51 | Penta_E | Penta_D | D8S1179 | FGA |
|---|---|---|---|---|---|---|---|
| C1R | 16<br>19 | 27<br>29<br>30 | 11<br>12 | 11<br>13 | 10<br>11 | 9<br>13<br>14 | 20<br>21<br>22<br>23 |
| C2S8 | 15<br>15 | 30<br>30.2 | 13<br>13 | 10<br>10 | 9<br>15 | 13<br>13 | 24<br>24 |
| C3H6 | 12<br>18<br>19 | 29<br>30 | 17<br>17 | 13<br>14 | 9<br>13 | 11<br>12<br>14 | 18<br>23 |
| C4C5 | 16<br>16 | 30.2<br>33.2 | 18<br>18 | 13<br>15 | 9<br>11 | 9<br>14 | 21<br>23 |
| C5H9 | 15<br>17 | 29<br>30 | 13<br>13 | 14<br>16 | 11<br>13 | 10<br>16 | 20<br>22 |

UAGC Staff
Approver Signature:

Stacy Sotak
