## Supplementary material for "DDB2 localizes to the nucleolus to mediate oxaliplatin resistance in colorectal cancer": Western blot full scan

### Slide 1
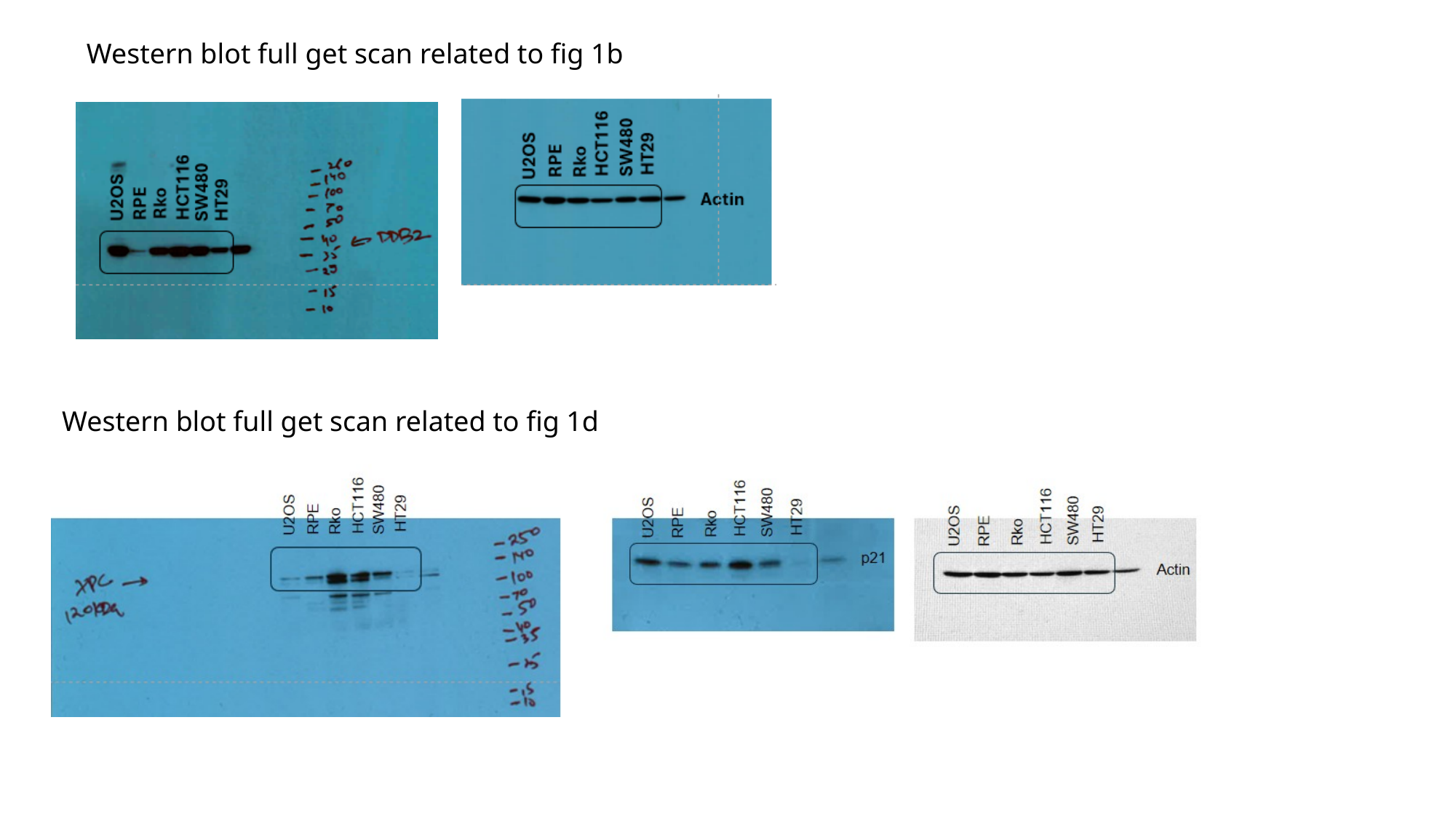

Western blot full get scan related to fig 1b
Western blot full get scan related to fig 1d

### Slide 2
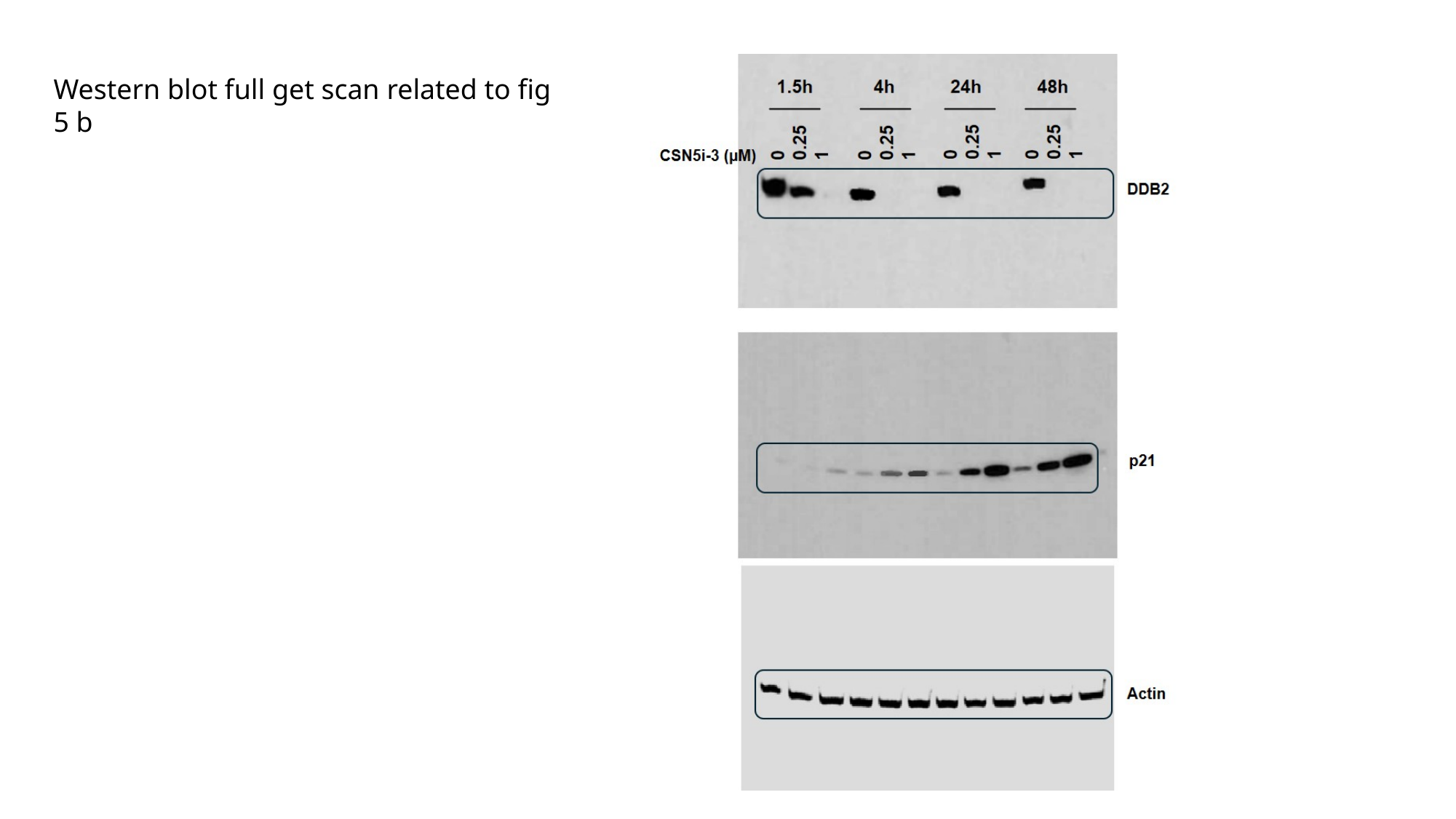

Western blot full get scan related to fig 5 b
